# Universal gut microbial relationships in the gut microbiome of wild baboons

**DOI:** 10.1101/2022.08.20.504530

**Authors:** Kimberly E. Roche, Johannes R. Björk, Mauna R. Dasari, Laura Grieneisen, David Jansen, Trevor J. Gould, Laurence R. Gesquiere, Luis B. Barreiro, Susan C. Alberts, Ran Blekhman, Jack A. Gilbert, Jenny Tung, Sayan Mukherjee, Elizabeth A. Archie

## Abstract

Ecological relationships between bacteria mediate the services that gut microbiomes provide to their hosts. Knowing the overall direction and strength of these relationships within hosts, and their generalizability across hosts, is essential to learn how microbial ecology scales up to affect microbiome assembly, dynamics, and host health. Here we gain insight into these patterns by inferring thousands of correlations in bacterial abundance between pairs of gut microbiome taxa from extensive time series data (5,534 microbiome profiles from 56 wild baboon hosts over a 13-year period). We model these time series using a statistically robust, multinomial logistic-normal modeling framework and test the degree to which bacterial abundance correlations are consistent across hosts (i.e., “universal”) or individualized to each host. We also compare these patterns to two publicly available human data sets. We find that baboon gut microbial relationships are largely universal: correlation patterns within each baboon host reflect a mixture of idiosyncratic and shared patterns, but the shared pattern dominates by almost 2-fold. Surprisingly, the strongest and most consistently correlated bacterial pairs across hosts were overwhelmingly positively correlated and typically belonged to the same family—a 3-fold enrichment compared to pairs drawn from the data set as a whole. The bias towards universal, positive bacterial correlations was also apparent in monthly samples from human infants, and bacterial families that had universal relationships in baboons also tended to be universal in human infants. Together, our results advance our understanding of the relationships that shape gut microbial ecosystems, with implications for microbiome personalization, community assembly and stability, and the feasibility of microbiome interventions to improve host health.

## Introduction

Mammalian gut microbiomes are highly diverse, dynamic communities whose members exhibit the full spectrum of ecological relationships, from strong mutualisms like syntrophy and cross-feeding, to competition, parasitism, and predation [1–4]. These relationships mediate a variety of biological processes that have powerful effects on host health and fitness, including the metabolism of complex carbohydrates and toxins, and the synthesis of physiologically important compounds, like short-chain fatty acids, neurotransmitters, and vitamins [1–8]. Despite their importance, major gaps remain in our understanding of microbial relationships in the gut microbiome [1, 9, 10]. We typically do not know if the abundance of one microbe consistently predicts the abundance of other microbes in the same host community, nor do we understand whether these correlative relationships are consistent in strength or direction across hosts ([10–13]).

Knowing the overall direction and strength of these correlative relationships is important, not only because they partly reflect the ecological relationships that mediate gut microbial processes, but also because overall correlation patterns can affect gut microbiome assembly, stability, and productivity [14, 15]. For instance, sets of microbes that exhibit strong, positive relationships within hosts sometimes represent networks of cooperating taxa that promote each other’s growth [5, 9, 16]. In turn, these strong, mutualistic interdependencies can create an ecological house of cards where microbes rise and fall together, hampering community assembly and stability [14, 17]. In addition, understanding the degree to which correlative relationships between microbes are the same or different in different hosts can shed light on whether hosts share similar, underlying microbial ecologies [9, 10, 18–20]. Filling this knowledge gap has consequences for the generalizability of microbiome assembly processes, stability, and the ecosystem services that emerge from microbiome dynamics to affect host health [9, 10, 12, 14, 17, 21].

To date, there are several reasons to think that correlative relationships in the gut microbiome will not be consistent across hosts and will instead be individualized to each host. For instance, several common community and evolutionary processes—such as horizontal gene transfer, genotype by environment interactions, and priority effects—can lead microbiome taxa to fill different ecological roles in different hosts [3, 22–26]. Further, some microbes can adopt context-dependent metabolisms and ecological roles depending on their microbial neighbors or other aspects of the environment—all phenomena that could lead to personalized interspecies relationships in gut microbiota [27–30]. Finally, the common observation that gut microbial community compositions (i.e., the presence and abundance of taxa) are highly individualized is sometimes proposed to arise from host-specific microbial ecologies and relationships [22–26, 31–35].

However, to date, the handful of studies that have tested the generalizability of gut microbial relationships across hosts suggest that these relationships are not highly individualized and are instead largely consistent (i.e., “universal”) across hosts (**Fig. 1A**; [10, 18–20, 36]). For instance, Bashan et al. [10] inferred “universal” gut microbial relationships in the human gut microbiome by applying dissimilarity-overlap analysis (DOA) to cross-sectional samples from several human data sets. DOA infers universal microbial relationships by testing whether pairs of hosts who share many of the same microbiome taxa also tend to have similar abundances of those taxa [10, 18–20, 36]. This approach relies on the assumption that, when two communities share many of the same species and have similar abundances of those species, they do so because of a shared, underlying set of between-species abundance relationships [10, 36]. While many studies using this approach find evidence that microbial relationships are “universal” [10, 18–20], DOA’s assumptions have been questioned because some conditions can lead to the spurious detection of universality, including environmental gradients, the strength of stochastic processes, and the presence of many non-interactive species [10, 36, 37].

**Figure 1.**
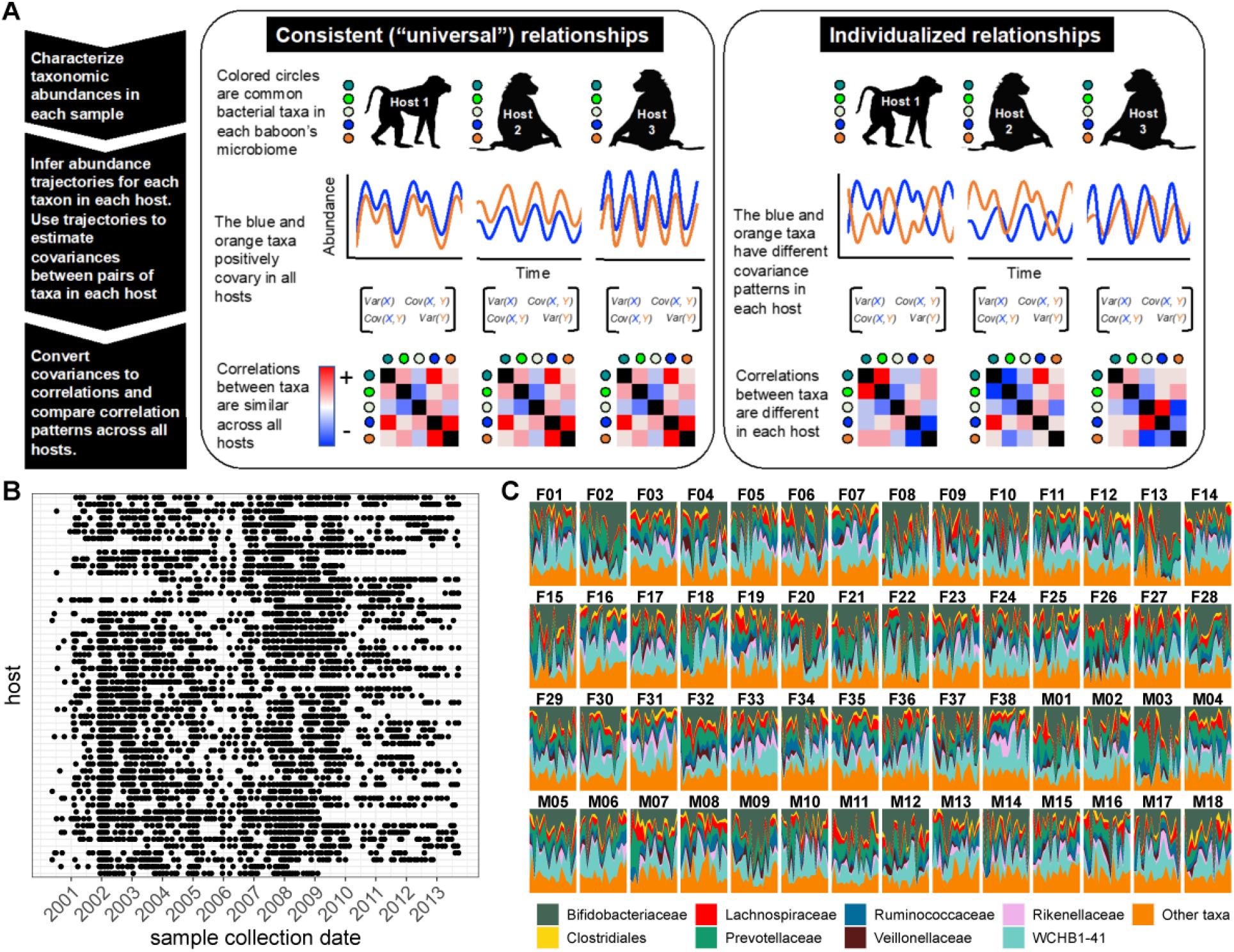
Testing the generalizability of gut microbial correlations across hosts. **(A)** Schematic illustrating our approach for testing the degree to which gut microbial abundance correlations are consistent (i.e., “universal” [10]) across different baboon hosts. The left-hand set of images show our expectations for consistent correlation patterns; the right-hand images show our expectation for individualized correlation patterns. Colored circles next to each baboon represent prevalent microbial taxa found in at least 20% of samples in each host (and excluding putative duplicate 16S gene copies; see methods). In each host, we inferred centered log-ratio (CLR) abundance trajectories for these taxa using a multinomial-logistic normal modeling approach implemented in the R package ‘fido’ [46]. Cartoons of two such trajectories for the orange and blue taxa are below each baboon. We used these trajectories to infer covariances between each pair of taxa in all baboons (represented by covariance matrices). We then converted these covariances to Pearson’s correlations and compared bacterial correlation patterns across all hosts, shown as heat maps (red cells are positively correlated taxa; blue cells reflect negatively correlated taxa). **(B)** Irregular time series of fecal samples used to infer microbial CLR abundance trajectories in 56 baboon hosts (n=5,534 total samples; 75-181 samples per baboon across 6 to 13.3 years). Each point represents a fecal sample collected from a known individual baboon (y-axis) on a given date (x-axis). Samples from the same baboon were collected a median of 20 days apart (range=0 to 723 days; 25th percentile=7 days, 75th percentile =49 days). **(C)** Relative abundances of the 8 most prevalent gut bacterial orders and families over time (x-axis) for all 56 hosts (samples from females are labeled with an F; male samples with an M). Microbiota were somewhat individualized to each host (**Fig. S2**; [42, 43]).

An obvious alternative is to measure microbial correlations directly from microbiome time series collected from several hosts [9, 38]. Unlike DOA, this approach should be able to pinpoint which microbiome taxa exhibit the most and least consistent relationships across hosts. However, measuring microbial correlations from longitudinal, multi-host microbiome time series has its own challenges: time series with adequately dense sampling are rare, and most such data sets exhibit temporal autocorrelation and irregular sampling [38]. Further, the most common, and still most feasible, way to collect microbiome community data—via high-throughput sequencing—generates noisy count data that usually can only be interpreted in terms of relative (not absolute) abundances [39, 40].

To overcome these hurdles, here we combine extensive time-series data on the stool-associated microbiota with a multinomial logistic-normal modeling framework (**Fig. 1**; n=5,534 samples from 56 baboons; 75 to 181 samples per baboon across 6 to 13.3 years, between 2000 and 2013; [41–43]). This framework uses 16S rRNA sequencing count data to learn a smoothly evolving Gaussian process. The baboon hosts were the subject of long-term research on individually recognized animals by the Amboseli Baboon Research Project in Kenya, which has been studying baboon ecology and behavior in the Amboseli ecosystem since 1971 [41]. The baboons range over the same habitat and experience similar diets and sources of microbial colonization, facilitating inference about the consistency of microbial correlations across hosts (**Fig. S1**; [42, 43]). Our modeling approach accounts for variation attributable to seasonal changes in the animals’ diets, proportionality in the count data, and irregularity in sampling to produce per-individual, per-taxon trajectories of log-ratio abundances that we used to estimate pairwise microbial correlations within each host.

We pursued four main objectives. First, we characterized the overall sign and strength of pairwise correlations in bacterial abundance within each host. Second, we tested the degree to which these correlation patterns are systematically consistent across hosts or individualized by host (**Fig. 1A**). Third, we identified taxonomic, phylogenetic, environmental, and host-related predictors of the direction and universality of bacterial correlations. Finally, we tested the generalizability of our findings by comparing the patterns of universality in our data set to two microbiome time series from humans [34, 44].

Our predictions for these analyses were influenced by ideas from community and microbial ecology. First, because strong interdependencies can hamper community assembly and destabilize community dynamics [14, 15, 17], we expected that most microbial correlations would be weak with few strong positive relationships between microbes. Second, consistent with studies that used DOA, we expected that microbial relationships would be more consistent across hosts than individualized (see **Fig. 1A** for a visualization of this prediction). This result would suggest that personalized microbiota—their compositions and dynamics—do not arise from host-specific microbiome ecologies [10, 18–20]. Third, because closely related gut bacteria may have similar functional properties, we expected to observe many positive correlations between those that are close phylogenetic relatives. Alternatively, competitive exclusion may lead closely related taxa to exhibit neutral or negative relationships. Fourth, because the environments experienced by baboons in Amboseli are far more uniform than those experienced by typical human study subjects [42, 43], we expected that the signature of “universality” in baboons would be stronger than that observed in humans. We discuss the implications of these patterns for individual microbiome community assembly and dynamics, and for understanding how microbiome communities are structured across hosts—a key requirement for successful intervention to improve host health [10, 11, 45].

## Results

### Most bacterial correlations within individuals are weak and negative

We began by characterizing the overall sign, strength, and significance of pairwise correlations in bacterial abundance within each host. To do so, we applied the approach outlined in **Fig. 1A** to stool-associated time series from 56 baboons (**Fig. 1B**) and calculated Pearson’s correlations between all possible pairs of bacterial taxa for three taxonomic partitions of the data. These partitions were: (1) all pairs of CLR-transformed amplicon sequence variants (ASVs) found in >20% of samples in each host and were unlikely to represent a duplicate 16S rRNA gene copy ([47]; see Methods; n=125 ASVs; **Fig. 2A; Table S1**); (2) all pairs of bacterial phyla found in >20% of samples in each host (n=12 phyla; **Table S2; Fig. S3**); and (3) all pairs of taxa agglomerated to the most granular possible family, order, or class found in >20% of samples in each host (n=37 taxa; **Table S3; Fig. S3**). We assessed the false discovery rate for each correlation with a threshold for significance of FDR ≤ 0.05, by comparing the nominal p-values for each observed correlation to an empirical permutation-based null, obtained by shuffling taxonomic identities within microbiome samples 10 times for each host and re-calculating the Pearson correlation p-values obtained from the permutations (**Fig. 2B**). We also confirmed that the resulting correlation patterns were insensitive to several modeling choices and were not primarily driven by seasonal shifts in microbiome composition (see results below and the Supplement).

**Figure 2.**
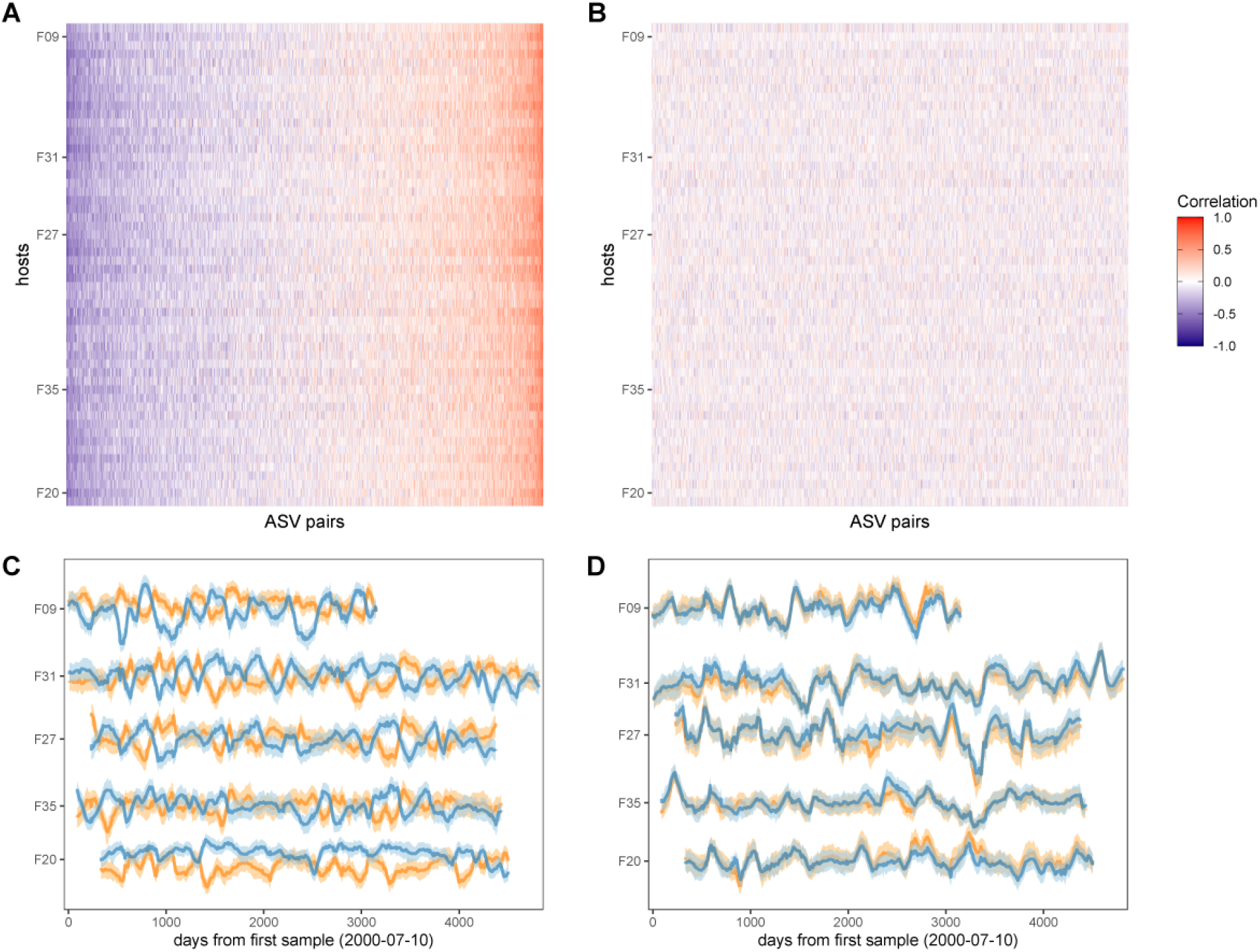
Bacterial correlation patterns across hosts. The heat map in panel **(A)** shows Pearson’s correlation coefficients of CLR abundances between all pairs of ASVs (x-axis) in each of the 56 baboon hosts (y-axis). Each pair of ASVs is represented on the x-axis, including all pairwise combinations of 125 ASVs resulting in 7,750 ASV-ASV pairs in each host (434,000 total correlations across all 56 hosts). Columns are ordered by the mean correlation coefficient between ASV-ASV pairs, from negative (blue) to positive (red). **(B)** Pairwise correlations generated from random permutations of the data. Taxonomic identities were shuffled within samples and pairwise ASV-ASV correlations were estimated to produce a null model of ASV-ASV correlation patterns within and between hosts. Column order is the same as in Panel A. Panels **(C)** and **(D)** show example trajectories of CLR abundances for two pairs of ASVs in the same five hosts. Panel **(C)** shows a strongly negatively correlated pair (median r across all hosts=-0.562; two ASVs in family Lachnospiraceae: ASV25 (orange) and ASV107 (blue); **Tables S1 and S4**) and panel **(D)** shows one strongly positively correlated pair (median r across all hosts=0.801; two ASVs in the genus Prevotella 9; ASV2 (orange) and ASV3 (blue); **Tables S1 and S4**).

Consistent with the expectation that most bacterial correlations in the gut microbiome are weak [14, 17], only 17% of ASV-ASV correlations in the heat map in **Fig. 2A** were stronger than expected by chance (FDR ≤ 0.05; **Fig. S4A**; 20% of phylum-phylum; 21% of family/order/class correlations; **Fig. S3**). The strongest negatively correlated pair in **Fig. 2A** included two ASVs in the family Lachnospiraceae that had a median correlation of −0.562 (+/- 0.118 s.d.) across all baboon hosts (**Fig. 2C**; ASV25 and ASV107; **Tables S1** and **S4**). The strongest positively correlated pair of ASVs included two members of the genus *Prevotella* that had a median correlation of 0.801 (+/- 0.053 s.d.) across all baboons (**Fig. 2D**; ASV2 and ASV3; **Tables S1** and **S4**). While these two ASVs were assigned to the same genus, their V4 16S DNA sequence identity was 97.6%, indicating they are probably not duplicate 16S gene copies in the same taxa [47] (**Table S4**).

In support of the idea that positive bacterial interdependencies are rare [14, 15, 17], only 8.8% of ASV pairs were significantly positively correlated within hosts over time, and the overall bacterial correlation patterns were slightly skewed towards negative relationships—especially for relationships between bacterial phyla. For instance, at the ASV-level, the median correlation coefficient in **Fig. 2A** was −0.016, and 53% of these correlations were negative (binomial test p < 0.0001). For family/order/class-level taxa, 55% of all correlations in were negative (**Figs. S3A** and **S4A**; median family/order/class-level correlation=-0.031; binomial test p < 0.0001). Correlations between phyla exhibited the strongest negative skew, with 64% of phyla-phyla correlations having a negative sign (**Figs. S3B and S4A**; median phyla-level correlation=-0.092; binomial test p < 0.0001). This bias towards negative relationships may reflect the fact that different phyla exhibit substantial differences in metabolism and lifestyle and likely respond to distinct environmental drivers.

### Within-host bacterial correlation patterns are largely consistent across baboons

Next, we tested the degree to which within-host ASV-ASV correlations were consistent across hosts. We began by plotting the absolute value of each ASV pair’s median Pearson’s correlation coefficient as a function of the consistency of their correlation sign (positive or negative) across the 56 hosts (**Figs. 3A and 3B**). These plots provide two main insights into the consistency of bacterial associations. First, in support of the idea that ASVs do not exhibit vastly different correlative relationships in different hosts, no ASV pairs were both strongly and inconsistently correlated across hosts (**Figs. 3A and 3B**; **Fig. S5A**). Instead, the ASV pairs that had inconsistent correlation signs across hosts always had weak and often non-significant median absolute correlation coefficients within hosts (**Figs. 3A and 3B**). Second, the pairs with the most consistent sign agreement across hosts also exhibited the largest median absolute correlation coefficients across hosts (**Figs. 3A and 3B**; Spearman’s r=0.844, p<0.0001). Hence, pairs of ASVs that have the strongest relationships, and are therefore likely to play the most important roles in structuring gut microbiome dynamics, also tend to have the most consistent relationships in different hosts. Indeed, for the sets of positively or negatively correlated ASV-pairs that showed universal agreement in the sign of their correlation across all hosts (i.e., where x=1 in **Figs. 3A and 3B**), the median correlation coefficient is 0.398, compared to 0.113 for those with no sign consistency (x=0.5 in **Figs. 3A and 3B**). Note, that the correlation strength for a given pair of ASVs was only weakly predicted by bacterial abundance. When both members of the pair were relatively abundant, pairs tended to exhibit stronger median correlations (r=0.012, p<0.0001; **Fig. S6**). However, while this effect is significant, it explained <1% of the variance in median correlation strength.

**Figure 3.**
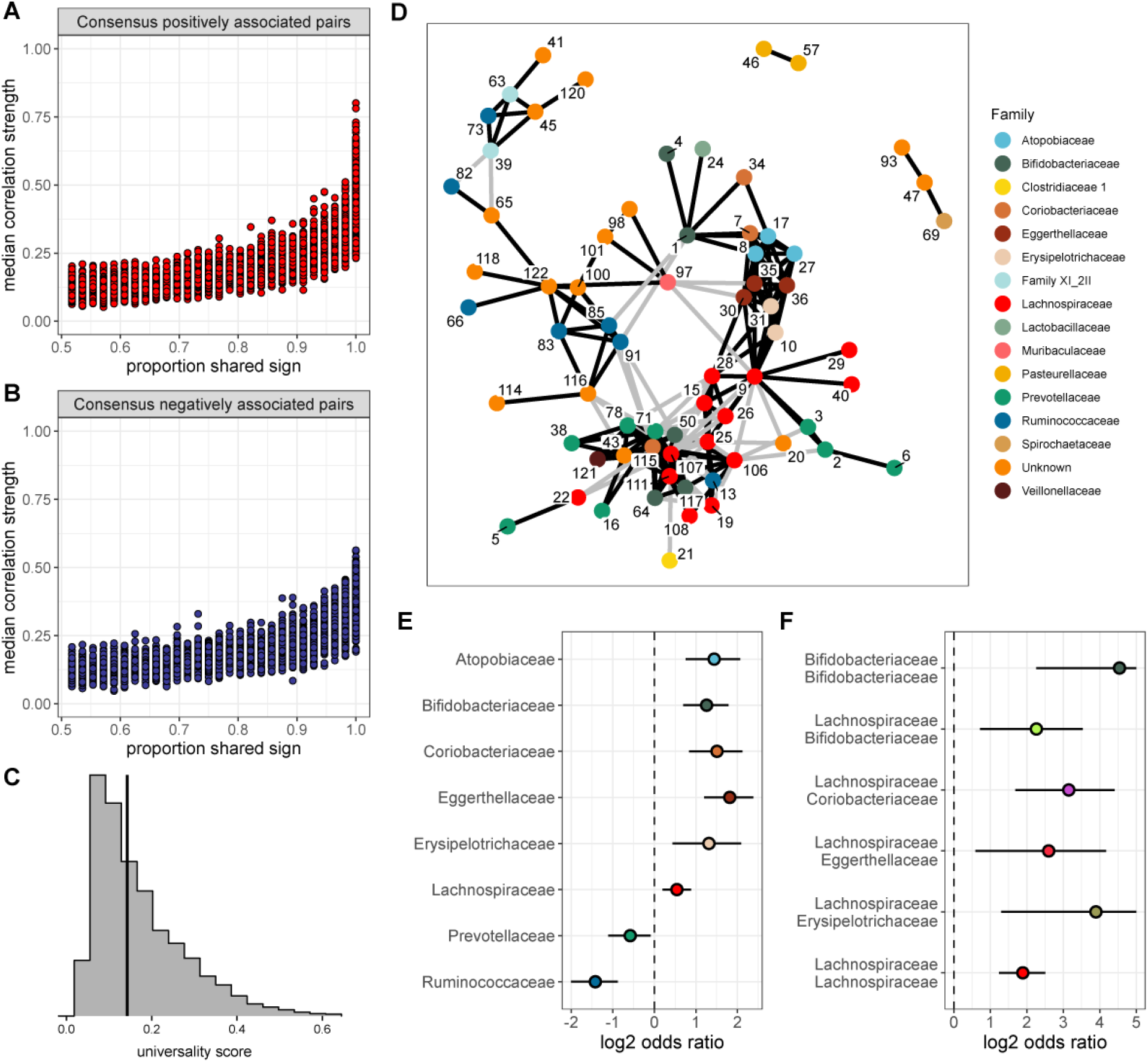
None of the ASV pairs were strongly and inconsistently correlated across hosts, and the strongest and most consistently correlated ASVs are typically positively correlated. Plots in **(A)** and **(B)** show the median correlation strength for each ASV-ASV pair across all 56 hosts as a function of the consistency in direction of that pair’s correlation across hosts, measured as the proportion of hosts that shared the majority correlation sign (positive or negative; ASV pairs that were positively correlated in half of the 56 hosts have a consistency of 0.5; ASV pairs that were positively [or negatively] correlated in all hosts have a consistency of 1.0). Panel **(A)** presents this relationship for consensus positively correlated features and panel **(B)** shows consensus negatively correlated features. The Spearman correlation between median association strength and the proportion of shared sign for all correlated features is 0.844 (p < 0.0001). Multiplying the two axes in either panel **(A)** or **(B)** creates a “universality score”, whose distribution is shown in panel **(C)**. This score reflects the strength and consistency of pairwise microbial correlations across hosts and ranges from 0 to 1, where a score of 1 indicates ASV-ASV pairs with perfect correlations of the same sign in all hosts. A vertical line indicates the minimum significant universality score. **(D)** Correlation networks for the top 2.5% most strongly and consistently correlated ASV pairs across hosts (i.e., the top 2.5% highest universality scores; pairs with rank 1-194 in **Table S4**). Network edges are colored by the consensus sign of the correlation between that pair (black for pairs where most hosts had a positive correlation; gray for pairs where most hosts had a negative correlation). Node labels indicate the ASV identity in **Table S1** and colors represent bacterial families. **(E)** Significantly enriched bacterial families in the network in panel D (Fisher’s Exact Test p < 0.01 all, FDR ≤ 0.05; see **Table S5** enrichment statistics for all families). **(F)** Significantly enriched same-family pairings in the network in panel D (**Table S5**). Note that for visualization, the estimated log2 odds ratio intervals have been truncated at 5; full estimates are given in **Table S5.**

Visual inspection of the patterns in **Figs. 2A**, **3A, and 3B** indicate that ASV-ASV correlations are largely consistent across baboons, as opposed to individualized to each baboon. To explicitly quantify the relative strength of shared versus individualized signatures in the heat map in **Fig. 2A**, we calculated the population mean pattern for the ASV-ASV correlation matrix, *m*. For each host, we then estimated the residual difference, *e*, between that individual’s observed ASV-ASV correlation matrix, *y*, and the population mean matrix: *y* – *m* (see **Fig. S7A** for a cartoon example). We reasoned that the observed correlation matrix for each host can be approximated by a mixture of contributions from the population mean matrix *m* and the host-specific residual matrix *e*. To identify the optimal mixture for each host (i.e., the mixture of consistent vs. individualized correlation patterns that best explained the observed data), we titrated the contribution (i.e., weight) of *e* from 0% to 100% (and correspondingly, the contribution of *m* from 100% to 0%) and identified the value that minimized the Frobenius distance between the simulated combination and the observed correlation matrix, *y*.

In support of prior observations of “universality” [10, 18–20], we found that, across hosts, the optimal mixture involved contributions from the shared correlation structure (i.e., *m*) of between 50% and 70% (median 65%) and a host-level contribution (i.e., from *e*) of between 50% and 30% (median 35%). Hence, population-level signatures contributed almost twice the weight as host-level signatures (a median population:host ratio of 1.86:1; **Fig. S7B**). As a result, ASV-level relationships tend to be more consistent across hosts than host-specific.

### The most consistent ASV-level correlations are positive and between phylogenetically related taxa

One advantage of our approach, compared to dissimilarity overlap analyses [10], is we can identify the bacterial pairs that exhibit the most consistent relationships across hosts. Hence, we next conducted several analyses to understand why some pairs of ASVs exhibit more consistent correlation patterns across hosts than others. To do so, we created a “universality” score that could be calculated for each ASV pair. The score multiplies the pair’s median correlation coefficient across hosts (y-axis of **Fig. 3A, 3B**) with its correlation consistency across hosts (i.e., proportion of shared sign; x-axis of **Fig. 3A, 3B**). The resulting scores range from 0 to 1, where a score of 1 equates to perfect “universality” (i.e., all hosts have a correlation coefficient of 1 or all hosts have a correlation coefficient of −1). Applying this score to all pairs of ASVs reveals a right-skewed distribution, reflecting the fact that most bacterial correlations are weak, with inconsistent sign directions across hosts (**Fig. 3C**; **Fig. S4B**). However, 49% of these scores were higher than expected by chance (permutation test; FDR ≤ 0.05; **Fig. 3C**; **Fig. S4B**), reflecting a signal of universality in our data.

Interestingly, the ASV-pairs with the highest universality scores almost always exhibited net positive correlations across hosts, as opposed to net negative relationships, suggesting that the most universal relationships occur between pairs of ASVs that respond similarly to shared drivers or facilitate each other’s growth. For example, among the ASV pairs in the top 1% of universality scores (n=78 pairs), 96.2% exhibited net positive correlations across all hosts, while only 5.6% (3 of 78 pairs) exhibited net negative correlations (**Table S4**). In the top 2.5% most universal ASV pairs (n=194), 78.4% had net positive correlations across all hosts (**Table S4**).

To visualize these highly consistent positive correlations, we plotted bacterial co-abundance networks connecting the top 2.5% most universal ASV pairs (**Fig. 3C**). A handful of ASVs were highly connected within this network. The most connected ASV was ASV107 (family Lachnospiraceae; **Table S1**; **Table S4**), which was connected to 20 other ASVs. Ten other ASVs were connected to more than 10 other ASVs, including six other members of Lachnospiraceae (ASV9, ASV25, ASV30, ASV106, ASV107, and ASV111), two members of Coriobacteria (ASV115 in the family Coriobacteriaceae and ASV30 in the genus *Slackia*), one member of Bifidobacteriaceae (ASV50), and one member of Prevotellaceae (ASV71). The ASVs involved in these top 2.5% pairs were enriched for the families Atopobiaceae, Bifidobacteriaceae, Coriobacteriaceae, Eggerthellaceae, Erysipelotrichaceae, and Lachnospiraceae (**Fig. 3E; Table S5**; all Fisher’s Exact Test p-values < 0.01; FDR ≤ 0.05).

The network in **Fig. 3D** revealed clusters of positive connections, often between ASVs assigned to the same family (**Fig. 3F**). In fact, same-family pairs were enriched by >3-fold in the 2.5% most universal taxon pairs (52 pairs observed vs. 19 expected, p < 0.0001). The cluster of interconnected red nodes in **Fig. 3D** represents members of Lachnospiraceae, and Lachnospiraceae-Lachnospiraceae pairings were 3.7 times more common in this network than overall (30 pairs observed vs. 9 pairs expected **Fig. 3F**). Bifidobacteriaceae also tended to exhibit within-family ASV pairings (**Fig. 3F**).

The observation that the most consistent correlations often occur among ASVs in the same family raises another question: does the phylogenetic distance between a pair predict the nature of their relationship? In support of the idea that closely related ASVs respond similarly to the environment or facilitate each other’s growth, we found a significant relationship between the universality score of a given pair of ASVs and their phylogenetic distance (Pearson’s r for positively correlated pairs=-0.213; p < 0.0001; **Fig. 4A**). In contrast, negatively correlated ASV pairs exhibited a weak positive relationship between phylogenetic distance and universality such that closely related taxa tended to be less universal than more distantly related taxa (Pearson’s r=0.049; p=0.004; **Fig. 4B**). In other words, the strongest and most consistently *negatively* correlated taxa tend to be only distantly related. Positively correlated, closely related pairs were often members of the families Atopobiaceae, Eggerthellaceae, and Lachnospiraceae, especially pairs where both members belonged to the family Lachnospiraceae (**Fig. 4C-D**; **Table S6).**

**Figure 4.**
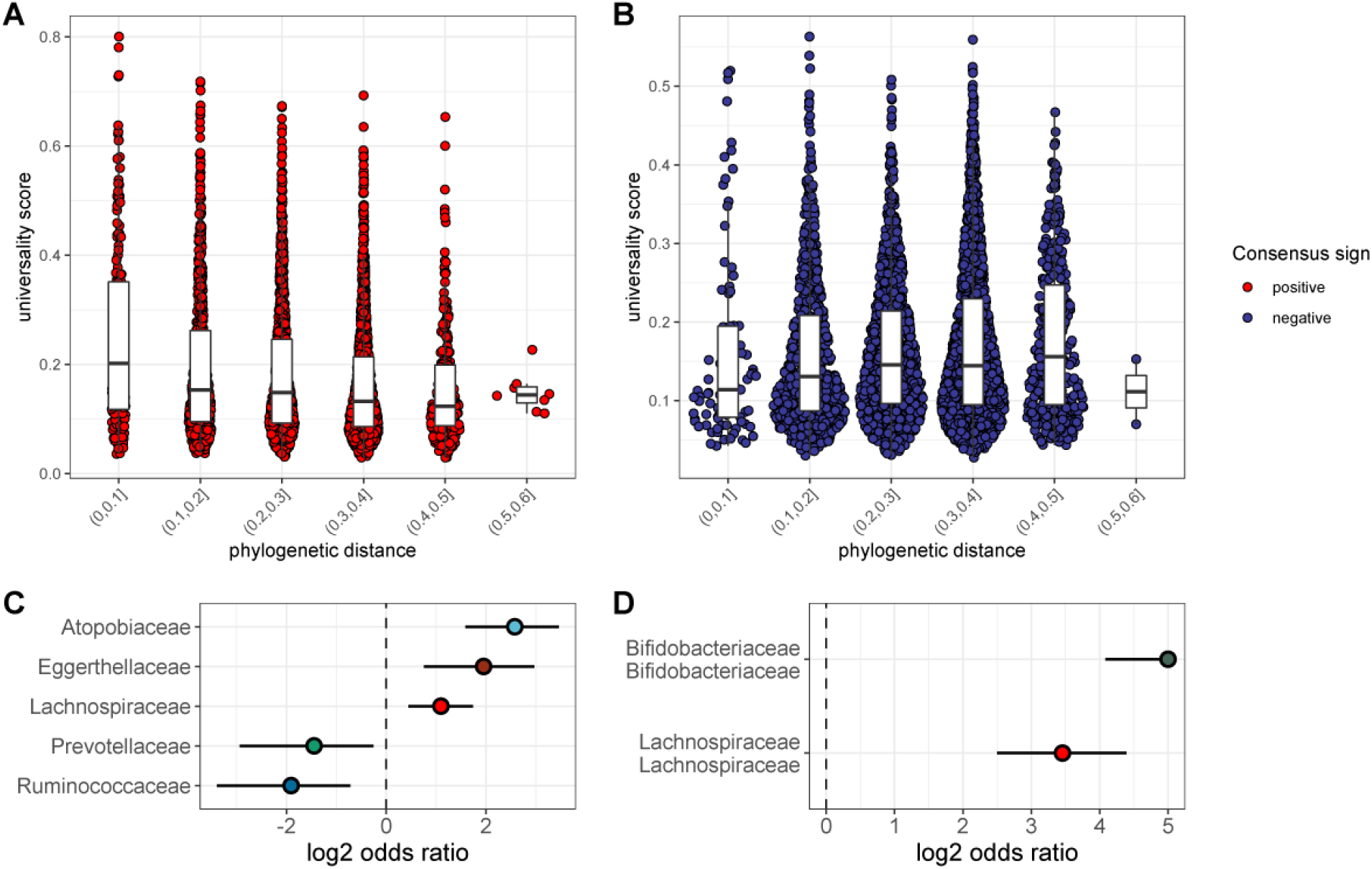
The most consistent ASV-level correlations are positive and often between close evolutionary relatives. Pairwise universality scores are plotted as a function of phylogenetic distance between the ASV-ASV pair for consensus positively correlated pairs in red **(A)** and negatively correlated pairs in blue **(B)**. Phylogenetic distance (x-axis) is binned into 0.1 increments; each point represents a given ASV pair, and box plots represent the median and interquartile ranges for a given interval of phylogenetic distance. Phylogenetic distance is negatively correlated with universality score in positive pairs (Pearson’s correlation for positively associated ASV pairs=-0.213, p-value < 0.0001), and positively correlated with universality score in negatively associated pairs (Pearson’s correlation for negatively associated ASV pairs=0.049, p = 0.004). Panel **(C)** shows families for the ASV pairs enriched in the closest related (distance < 0.2) and highly universal (score > 0.5) pairs. Panel **(D)** shows enriched family-family pairings for the same subset of closely related and highly universal ASV pairs in panel C. Note that for visualization, the estimated log2 odds ratio intervals have been truncated at 5, which excludes 5 pairs with high uncertainty in the odds ratio; full estimates are given in **Table S6.**

### Genetic relatives, and hosts with similar microbiome compositions, have more similar bacterial correlation patterns

We next asked whether host-level variables, including sex, social group membership, genetic relationships, and baseline gut microbiome composition predict host differences in patterns of bacterial correlation. Consistent with prior research [10], the strongest predictor of distance in bacterial correlation patterns was distance in terms of baseline microbiome composition. Indeed, a Mantel test correlating compositional distance of average microbial profiles (as Aitchison distances between the per-host mean of centered log-ratio-transformed samples) with distance in microbial correlation patterns between hosts (via Frobenius distance) revealed that 34% of the variation in correlation patterns was explained by baseline microbiome community composition (Mantel: r^2^=0.343; p=0.001; **Fig. 5A**; **Table S7**).

**Figure 5.**
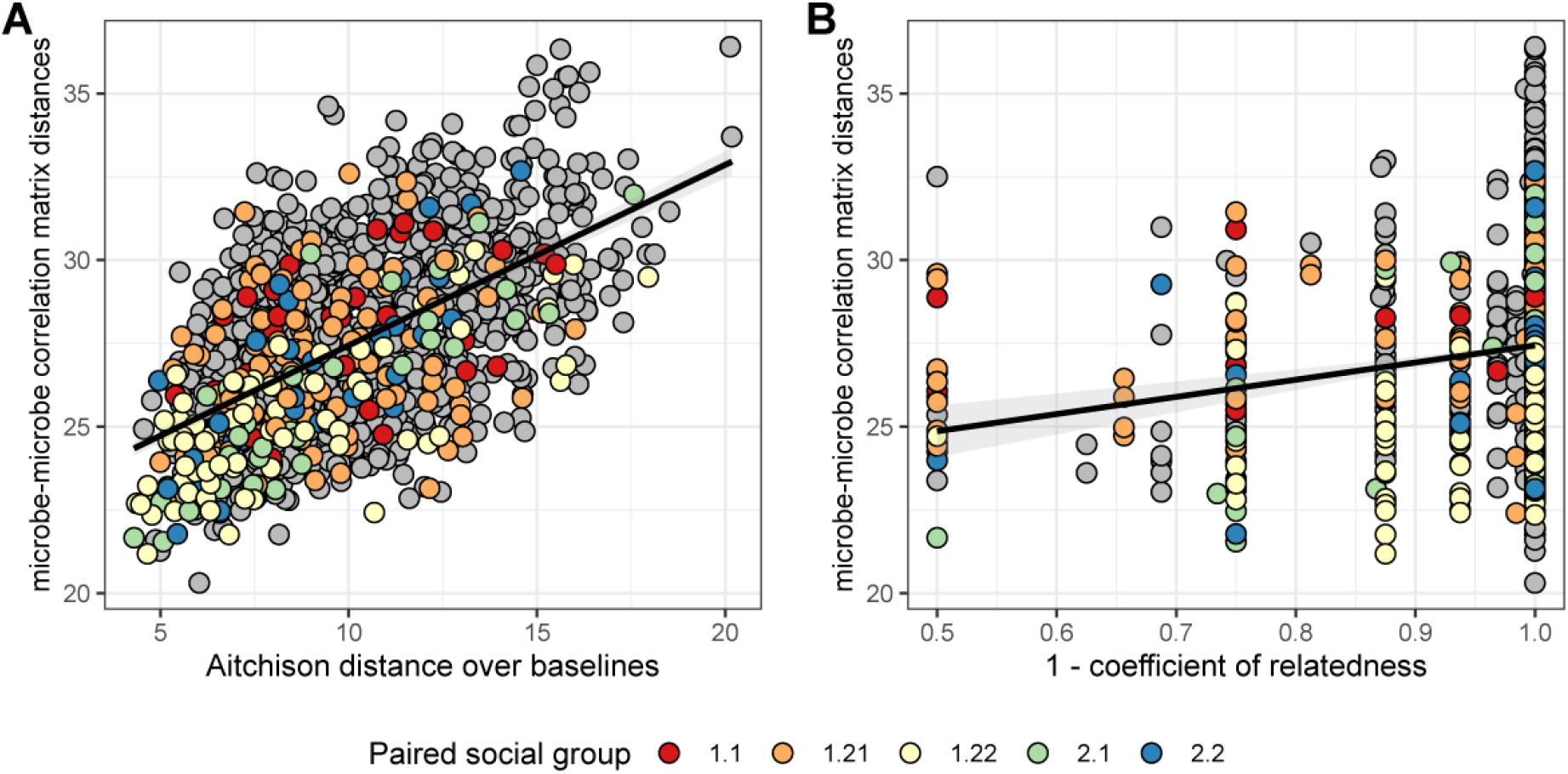
Baboons with more similar bacterial correlation patterns are more likely to have more similar baseline microbiome compositions and are more likely to be genetic relatives. In panel **(A)** each point is a pair of hosts; the y-axis shows the similarity of these hosts’ bacterial correlation patterns (via Frobenius distance) as a function of their microbiome compositional similarity (via Aitchison distance; Mantel: r^2^ =0.343; p=0.001). Colors show samples from pairs of baboons living in the same social group and grey dots are pairs of animals living in different social groups; there is no detectable effect of social group on correlation pattern similarity. Panel **(B)** shows the same Frobenius distances as a function of host genetic dissimilarity (1 – the coefficient of genetic relatedness between hosts; R^2^=0.025; p-value Mantel test 0.001). Colors reflect pairs of hosts living in the same social group, as in panel A.

Consistent with prior research in our population, which finds widespread heritability of the abundance of individual gut microbiome taxa [43], we also found a weak but significant relationship between host genetic distance and the distance in microbial correlation patterns between hosts. Hosts who were more distantly related based on a multigenerational pedigree have slightly less similar ASV-level correlation matrices, as measured by Frobenius distance (**Fig. 5B**; **Table S7**; r^2^=0.025; Mantel p-value=0.001). We found no evidence that members of the same social group or sex exhibit especially similar microbial correlation patterns (social group: F=1.994; p=0.106; sex: F=1.784; p=0.187; **Table S7**).

### Universality in Amboseli is not solely explained by seasonality or synchrony

Without experiments, we cannot disentangle whether our observed bacterial correlations are due to ecological interactions between bacterial species (e.g., mutualisms, direct or indirect competition etc.) or to shared responses to environmental gradients. While our modeling approach accounts for seasonal changes in the first three principal components of the baboons’ diets, to identify other potential effects of season we re-estimated the ASV-ASV correlation matrix after removing an oscillating seasonal trend from the observed log-ratio abundance for each ASV (**Fig. S8**). Removing this trend had little effect on the ASV-ASV correlation matrix; the variance explained by the seasonal oscillation is small for all ASVs (median 1.1%, minimum=0%, maximum=6%) and the between-ASV correlation estimates were almost identical to those derived from our original model (Pearson’s r=0.979, p<0.0001; **Fig. S8C**). We also tested whether pairs of ASVs with especially consistent between-host correlation patterns tend to show large seasonal changes in CLR abundance. To do so, we focused on 13 families that exhibit significant seasonal changes in CLR abundance, based on a previous analysis of the same data set [42]. While ASV pairs in which one member belongs to one of these significantly “seasonal” families are slightly more universal, this effect is small (difference of 0.026, p<0.0001 vs. pairs where 0 or 1 partner were “seasonal”; **Fig. S9**).

Because the high level of universality we observed was not well explained by season, we also tested whether universality is explained by synchronized dynamics. We reasoned that if one member of an ASV pair shows highly synchronized dynamics across different hosts, and the other member is also strongly synchronized across hosts, then universality could be an inevitable outcome of each member of the pair’s strong synchrony. We quantified synchrony as the degree to which the observed dynamics of a single, focal ASV are consistent across hosts, such that high synchrony (near 1) implies that the timing and direction of shifts in log-ratio ASV abundance are identical across hosts in the population (see Methods; **Fig. S10**). Estimates of synchrony ranged from 0.019 to 0.477 (median=0.196). Interestingly, ASVs in the 13 “seasonal” families are not more likely to have high synchrony than other families (ANOVA, p=0.358; **Fig. S11**). However, the average synchrony of an ASV-ASV pair did predict that pair’s universality score (r=0.264, p<0.0001): ASV pairs that are more synchronous on average are also more likely to show consistent correlations across hosts. These high synchrony, high universality pairs are highly enriched for Bifidobacteriaceae-Bifidobacteriaceae and Lachnospiraceae-Lachnospiraceae family pairs (**Fig. S12**).

### Baboon microbiomes are not substantially more “universal” than human microbiomes

Finally, to investigate the generalizability and applicability of our observations in baboons, we turned to two publicly available gut microbial time-series data sets: daily samples from 34 adults over a 17-day span (483 total samples; hereafter “Johnson et al.” [34]), and the DIABIMMUNE cohort that consists of 285 samples, collected monthly over 3 years, from 15 infants and toddlers living in Russian Karelia ([44]; at the time of writing, these cohorts were the only publicly available data sets we could find that included large numbers of repeated samples from the same subjects). Because baboons in Amboseli experience less heterogeneity in their environments and diets than humans [42, 43], we expected they would exhibit greater consistency in microbial correlations than either human cohort. Note that we compared each host cohort’s universality at the family/order/class level because this taxonomic level offered the greatest comparative power (10.1% of families/orders/classes overlap between the cohorts compared to just 3.1% of genera and no ASVs).

Contrary to our expectations, we find comparable evidence of universality in baboons and the DIABIMMUNE infant/toddler cohort, but weak universality in Johnson et al. (**Figs. 6A-6D**). Bacterial families in the DIABIMMUNE cohort yielded universality scores slightly higher than those observed in Amboseli (25th percentile=0.132, median=0.206, 75th percentile=0.316 for DIABIMMUNE; 25th percentile=0.088, median=0.150, 75th percentile=0.234 for Amboseli), driven by relationships between families that were stronger on average than those estimated for baboons (median DIABIMMUNE family-family correlation strength=0.270; median Amboseli family-family correlation strength=0.181). The high level of consistency between both infants/toddlers and baboons in one wild population is surprising and may point to the similar sampling intervals for these cohorts. Both cohorts were sampled approximately monthly, while Johnson et al.’s subjects were sampled daily [17, 48]. Median correlation strengths and universality scores for the Johnson et al. [34] cohort were substantially lower (median correlation=0.099; 25th percentile universality=0.050, median=0.076, 75th percentile=0.111) than the DIABIMMUNE cohort or the baboons.

**Figure 6.**
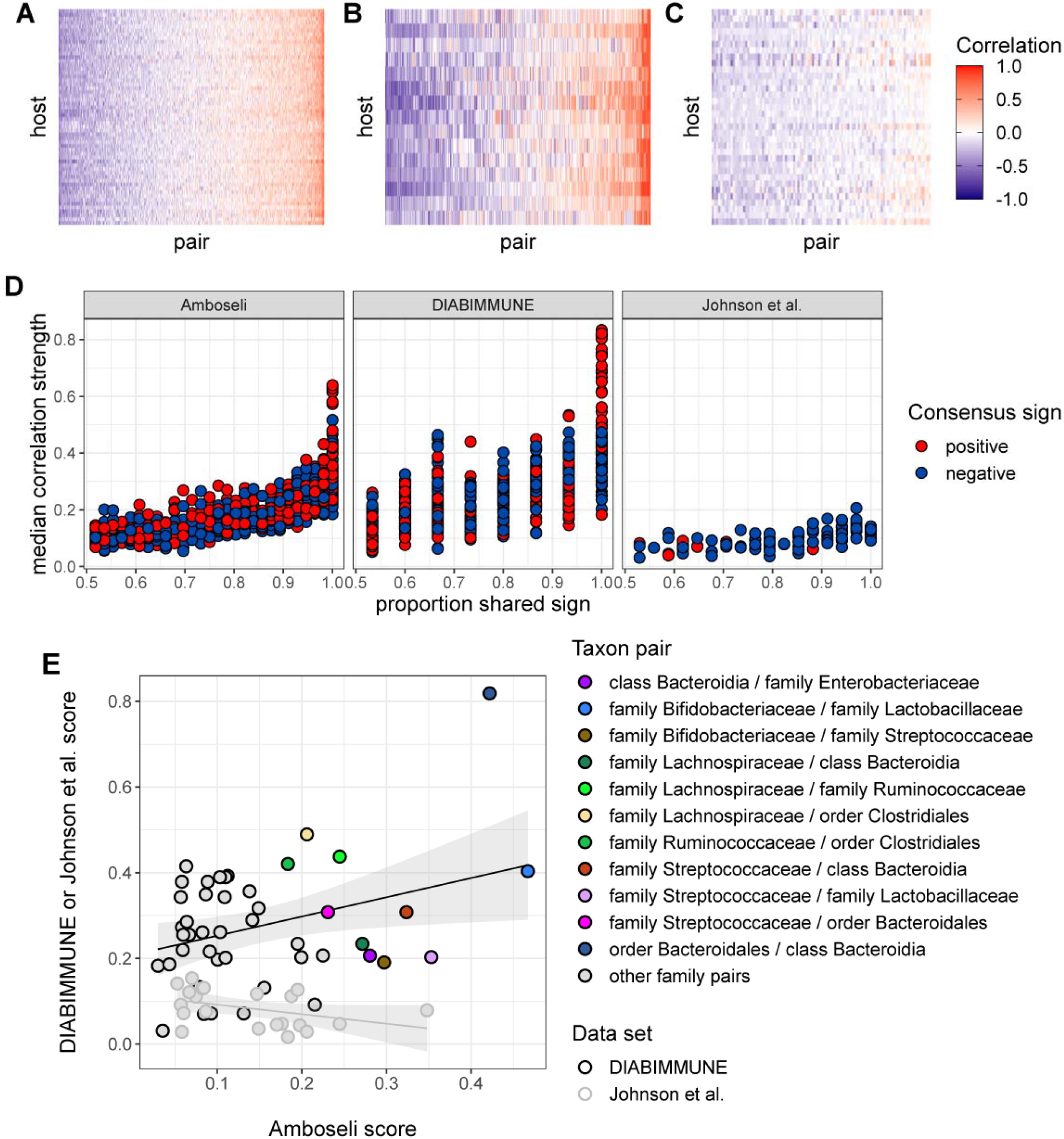
Patterns of universality in baboons are recapitulated in the DIABIMMUNE study. Following **Fig. 2A**, Panels **(A)**, **(B)**, and **(C)** show the Pearson’s correlation coefficients of CLR abundances between all pairs of families (x-axis) in two time series data sets from human subjects: **(A)** the Amboseli baboons, **(B)** the DIABIMMUNE cohort, consisting of 15 infants/toddlers sampled monthly over 3 years in Russian Karelia [44], and **(C)** the diet study of Johnson et al. [34], including 34 adults sampled daily over 17 days. Following **Figs. 3A and B**, panel **(D)** shows the median correlation strength of each family pair’s correlation coefficient across hosts as a function of the consistency in direction of that pair’s correlation across hosts (i.e., the proportion of hosts that shared the majority correlation sign, positive or negative). Median correlation strength is low overall in Johnson et al. (median=0.099), whereas the Amboseli baboon and DIABIMMUNE infant/toddler cohorts show similar relationships between median correlation strength and the proportion shared correlation sign across hosts (Spearman’s r in Amboseli=0.844; Spearman’s r in DIABIMMUNE=0.686). **(E)** Universality scores for overlapping family pairs from the infant/toddler subjects of the DIABIMMUNE study and baboons in the Amboseli study are significantly correlated (r=0.449, p=0.0226). Panel D shows universality scores for overlapping gut bacterial family pairs in the Amboseli baboon and DIABIMMUNE infant/toddler data sets (black outlines), as well as the Amboseli and Johnson et al. data sets (gray outlines) on opposing axes. Color represents the taxonomic identities of the family pairs.

Despite considerable differences in the hosts, time scales, and designs of these studies, all three data sets exhibited a positive correlation between correlation strength and sign consistency for family pairs (**Fig. 6C**). This correlation was strongest in the Amboseli baboons (Spearman’s r=0.844; p<0.0001); weaker in the DIABIMMUNE cohort (r=0.686; p<0.0001) and weakest in Johnson et al. [34] (r=0.644; p<0.0001). Further, the observation that the most universal family-family associations skew positive in baboons was replicated in the infant data set, but not in Johnson et al. [34]. All of the top 1% and top 2.5% most universal family pairs (6 of 6 and 16 of 16 pairs, respectively) are positively associated in the DIABIMMUNE cohort, compared to 86% and 71% of these pairs in the Amboseli baboons.

Finally, we examined the relationship between universality scores for family pairs that overlapped between Amboseli and DIABIMMUNE (n=45 pairs), and between Amboseli and Johnson et al. [34] (**Fig. 6D**; n=21 pairs; only 10 family pairs overlapped between all three data sets). For these overlapping pairs, scores in the Amboseli data predicted scores for the same family-family pair in the DIABIMMUNE data set (r=0.449, p=0.023). The association between scores in the Amboseli data and the Johnson et al. data was negative, but not statistically significant (r=-0.222, p=0.071).

## Discussion

Do different hosts have different microbiome “ecologies”? Answering this question is essential for predicting gut microbiome community assembly and dynamics, and for understanding the degree to which the species interactions that govern these processes are shared across hosts. Here, we overcome the constraints of previous cross-sectional analyses by measuring bacterial correlations directly from longitudinal, multi-host microbiome time series. Our results provide independent confirmation for prior studies that tested for universal gut microbial relationships via dissimilarity overlap analyses (DOA; [10, 18–20, 36]). We confirm that bacterial correlation patterns are largely shared across hosts in the same population, as opposed to idiosyncratic to individual hosts, and that hosts with the most similar bacterial correlation patterns are those with the most similar baseline microbiome compositions—a core assumption of DOA. Because prior analyses of these microbiome time series find that each baboon exhibits a highly personalized microbiome composition and dynamics [42], our findings suggest that such compositional personalization, which is also common in humans [22–26, 31–35], cannot be easily explained by personalized microbiome ecologies. Further, in terms of microbiome therapeutics, our results suggest that widely applicable microbiome interventions may be more attainable than personalized microbiome compositions would suggest.

By measuring bacterial correlations in multiple hosts, we were also able, for the first time, to pinpoint which pairs of bacterial taxa exhibit the most consistent relationships across hosts. Surprisingly, we found that the most universal bacterial pairs are almost always positively (as opposed to negatively) correlated. Positive bacterial interactions have been the subject of recent controversy [9, 15, 49]. Ecological theory predicts that strong positive interactions should be rare in natural communities because species interdependencies can hamper community assembly and stability [14, 17]. This theory is supported by experiments that directly measure the effects of one bacterial species on another’s growth [50–53] (but see [49]). Our results suggest that positive bacterial correlations are indeed uncommon in intact, unmanipulated microbiomes: significant positive relationships made up just 8.8% of all of the pairwise correlations we observed. However, when they occur, they often contain taxa that belong to the same bacterial families or are otherwise phylogenetically close, suggesting they may be members of the same ecological guild and respond similarly to shared resources and other environmental drivers. This pattern may partly explain the abundance of positively associated Lachnospiraceae pairs in our data, a family in which positive, within-family interactions are known to contribute to hydrolyzing starch and other complex carbohydrates, and ultimately the regulation of short chain fatty acids (SCFAs) [54–56].

These observations—that bacterial correlation patterns are largely consistent across hosts, and that the most consistent correlations are typically positive—were also apparent in one human data set, despite differences in study design, host age, and time scale. Specifically, both the Amboseli baboons and the DIABIMMUNE infant/toddler cohort from Russia [44] exhibit comparable levels of universality. This outcome was surprising, given that baboons are expected to experience less heterogeneity in their environments and diets than human children from birth to age three years—even if those infants are from the same population (Russian Karelia). We also found that the most universal bacterial families in baboons tended to be highly universal in human infants/toddlers. Hence, some bacterial families may exhibit consistent microbial relationships within hosts, across host populations, and across host species. Finally, a recent, independent study also identified consistent bacterial correlation patterns across four different populations of human hosts [9]. While this study lacked resolution at the level of individual hosts, it did identify a highly conserved network of positively associated and closely related microbes similar to those we identify in **Fig. 3**. The authors speculate that these conserved associations may indicate strong partner fidelity or obligate partnerships.

We did, however, fail to detect universality in a second human data set reported in Johnson et al. [34], in which subjects were sampled daily, rather than weekly or monthly. The lack of universality in Johnson et al. [34] may be due to this difference in sampling time scale, especially if daily abundances and correlations are noisier than covariances modeled over the longer time scales in our study. In support, many fewer of the microbial correlations were stronger than random chance in Johnson et al. as compared to the baboons or children in the DIABIMMUNE cohort. The subjects in Johnson et al. [34] also consumed substantially different diets from each other, perhaps more so than the children in the DIABIMMUNE cohort, and this inter-host difference in diet may reduce the universality of microbial correlations.

In terms of understanding microbiome ecology, an essential caveat to our findings is that the correlation patterns we observed could reflect either direct or indirect relationships, or uncontrolled environmental gradients, and they cannot be mapped directly to standard categories of pairwise ecological interactions, such as mutualism, commensalism, amensalism, exploitation, or competition. Experimental approaches that directly measure the effects of one species on another’s growth *in vitro* are better suited to characterizing these relationships [49–53]. However, even then, caution is required because a microbe’s community and environmental context can have important consequences for its metabolism, functional capacities, and relationships with other microbes. We surmise that most of the correlation patterns we observed are not attributable to environmental gradients because our signature of universality persisted, even when we accounted for diet, oscillating seasonal drivers, and microbial synchrony between hosts. Hence, some of correlations we observed may derive from microbial interactions themselves, rather than shared environmental drivers creating shared dynamics.

Our finding that correlations between gut microbial taxa are largely consistent across hosts is important, considering that many studies find highly personalized gut microbial compositions and single-taxon dynamics [27–29]. Personalized compositions and dynamics in the gut microbiome are commonly attributed to horizontal gene transfer and functional redundancy, which may lead some bacteria to perform different functions and exhibit different environmental responses in different hosts. Our results suggest these processes do not substantially alter pairwise microbial associations in the gut, at least for highly prevalent taxa at the level of ASVs and above, and on the time scales in our study (on the order of weeks and months). Because ASVs encompass multiple species and strains, reflecting the functional diversity of these taxa, their dynamics may be somewhat buffered against idiosyncrasies driven by horizontal gene transfer and functional redundancy, which affect single strains more strongly than whole species or genera. If so, personalized gut microbial compositions may emerge instead from personalized assembly processes [57, 58], the fact that most microbial relationships are weak, and the effects of rare, host-specific taxa (which were necessarily excluded from our analyses). A logical next step would be to confirm the stability of the microbial correlations we observed using culture-based approaches, which will help reveal the stability of these correlations in vitro and whether they can be attributed to direct effects of one microbe on another’s growth.

## Methods

### Study population and microbiome profiles

The baboon hosts in this study were members of the Amboseli baboon population, which has been studied by the Amboseli Baboon Research Project since 1971 [41]. The microbiome compositional profiles are derived from V4 16S rRNA gene amplicon sequencing data that were previously analyzed in [42, 43]. Our analyses use 5,534 of these profiles from 56 especially well-sampled baboons, collected over a 13.3-year span between 2000 to 2013 (**Fig. 1B**). Each baboon host in this data set was sampled at least 75 times (mean number of samples=99; range=75 to 181 samples; median number of days between samples within hosts=20 days; 25th percentile=7 days, 75th percentile =49 days). DNA was extracted from each sample using the MoBio and QIAGEN PowerSoil kit with a bead-beating step. All samples were sequenced on an Illumina HiSeq 2500, with a median read count of 48,827 reads per sample across all 5,534 samples (range=982 to 459,315 reads per sample). Further details of sample collection, DNA extraction, and sequencing can be found in [42, 43].

### Filtering of low-abundance taxa

Data sets of per-sample taxonomic counts were produced at each of three taxonomic levels, from finest to coarsest: ASV, taxonomic assignments finer than phyla, but above the genus level (e.g., class, order, family), and phylum. At the intermediate and coarsest levels, taxa were agglomerated using phyloseq’s tax_glom() function [59] such that all sequence variants sharing taxonomic identity at that level were collapsed into a single taxon (e.g. family Bifidobacteraceae).

To reduce sparsity in the data set, remove 16S sequences that could represent gene duplications, and focus only on taxa that were prevalent in all 56 hosts, we further filtered as follows: (1) in each of the three taxonomically defined data sets (i.e. ASV, taxa assigned to family/order/class, and phylum), we identified taxa present in a minimum of 20% of each host’s samples; (2) if a given ASV was >99% genetically similar to another ASV we removed the least abundant of the pair to minimize the risk of including duplicate 16S rRNA gene copies from the same taxa [47]; and (3) counts associated with all other taxa were combined into a dummy category, hereafter referred to as “other.” The “other” category therefore includes a combination of rare and host-specific gut microbes. This category was retained in the data set (although not analyzed directly) because “other” counts still inform the precision of the observed relative abundances in our model. Characteristics of the filtered data at each taxonomic level are provided in **Tables S1-S3**.

### Modeling log-ratio dynamics

Estimates of taxon-taxon covariance were obtained from the *basset* model of the “fido” package in R [46]. Data for each host took the form of a *D* × *N* count matrix, where *D* gives the number of taxa and *N* the number of samples collected for a given host. The following model was fit to each host’s count matrix (*Y*) where *Y_i_* represents the counts associated with a single sample:

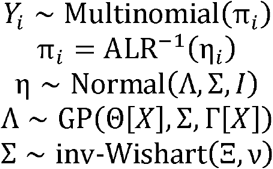

The observed relative abundances are considered to be drawn from a multinomial distribution parameterized by a set of proportions (π) which have an analogous representation in the additive log-ratio. The dynamics of these log-ratio abundances (η) are described by what amounts to a state space model in the 3^rd^ and 4^th^ lines of the specification above, where a Gaussian process models the evolution of a “latent” state. The matrix Σ captures covariation in log-ratio abundances (the *D* rows of the observed count matrix). Sample-sample covariation arising from nearness in time (autocorrelation) is modeled by the kernel matrix Γ. Both the kernel matrix and the expected baseline log-ratio abundances (Θ) are parameterized by a set of time-varying covariates *X* which included the day of sampling (where the date of first sample is defined as zero) and the first three principal components of diet composition, calculated following [42, 43] as the diet all juveniles and females living in the host’s social group in the 30 days prior to sample collection. All group members consume highly similar diets as they travel in a together across the habitat, encountering the same resources at the same time [42, 43]. These data are collected via random-order behavioral observations collected two to four times per week on adult females and juveniles in each social group. Parameterization of the kernel matrix is further described in the Supplement.

Posterior inference on this model is performed as described in [46] and yields estimates of the distributions of parameters necessary to reconstruct trajectories for all log-ratio taxa across sampling time. In particular, we extract the posterior estimates of one such parameter, Σ, the covariance of additive log-ratio (ALR) taxa, from the fitted models for each host. We convert these covariance matrices over ALR taxa to the centered log-ratio (CLR) form (a simple linear transformation of the matrix). We then normalize estimated CLR covariance matrices to Pearson correlation matrices in R using the built-in cov2cor() function.

### Calculating universality scores for taxon-taxon pairs

We devised a universality score for each pair of taxa intended to capture the strength and consistency of taxon-taxon correlations across hosts. The majority direction is negative otherwise. This score identifies the sign of the taxon-taxon correlation (positive or negative) that is most common across the 56 hosts (i.e., occurs in >50% of the 56 hosts in the data set). The direction of this sign is the “majority correlation sign.”

For a pair of taxa *i*, let 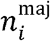 be the number of hosts with CLR correlation over pair *i* with the majority correlation sign for that pair and let *n* be the total number of hosts. Let *R* be the subset of estimated CLR correlations for pair *i* across hosts with the majority sign. The universality score *u_i_* for that taxon-taxon pair is then given by

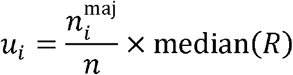

This score is the product of the median CLR correlation across hosts and the proportion of hosts with the majority correlation sign, and is bounded between 0 and 1. Scores near 1 indicate strong universality and near-zero scores indicate weak universality. Strong universality can only be achieved by taxon-taxon correlations that are both large in magnitude and highly concordant across hosts.

### Defining a cutoff for significant bacterial correlations and universality scores

We identified correlations stronger than expected under random simulations using permutations of the data set to define empirical null distributions (**Fig. S4A**). Specifically, we permuted the data by randomly shuffling taxon identity within each sample 10 times for each of the 56 hosts. This procedure maintained relative abundance patterns within a sample but scrambled the covariance patterns of relative abundances. The distributions of ASV-level CLR correlations in the original and permuted data are shown in **Fig. S4A**. We identified “significant” correlations as those below FDR ≤ 0.05 (Benjamini-Hochberg), testing against the permuted data.

We applied an analogous permutation test to derive a null distribution for taxon-taxon universality scores. In a single iteration of this permutation procedure, rows and columns of the observed taxon-taxon correlation matrix for each host were shuffled, maintaining the distribution over observed correlations at the host level but randomizing the identity of taxon pairs across hosts. This procedure was repeated 100 times and universality scores were calculated from each of these shuffled data sets to give a pseudo-null distribution of universality scores. The observed and null distributions of universality scores at the ASV level are shown in **Fig. S4B**. We used this empirical null distribution to identify universality scores significantly greater than expected (FDR ≤ 0.05).

### Estimating the ratio of population-level to host-level contributions to observed taxon-taxon correlation patterns

We used simulations to estimate the degree of shared “signal” between hosts in terms of taxon-taxon correlations. Each host’s “observed correlations” were defined as the *basset* estimated maximum a posteriori (MAP) estimates of centered log-ratio ASV correlations for that host. We computed the *mean* correlations across the population using the function estcov() from the shapes package in R [60] and estimated a host-specific contribution to the observed correlations as the residual *difference* between per-host observed and these mean correlations. That is,

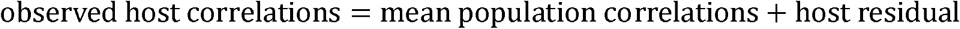

For each host, we simulated a hypothetical set of composite taxon-taxon correlations as a convex combination of mean and host residual:

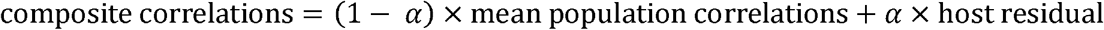

A cartoon example of this procedure is given in **Fig. S7A**. For example, one such simulated set of taxon-taxon correlations might constitute a mixture of 90% host contribution and 10% shared population-level “signal” (α=0.9). Alternatively, a small host-level contribution might have α=0.1.

For each host, we iterated over increasing proportions of host-level contribution (from 0% to 100%), generating simulated composite correlation matrices according to the formula above. We compared these simulated patterns to those observed for the same host, reasoning that simulated correlation matrices that minimize the distance between the observed correlation matrices and the simulated mixtures provide the best description of the underlying true mixture.

### Estimating synchrony

Seasonal autoregressive models were fit independently to each CLR-transformed ASV with arima() in R, using covariate matrices which included per-host intercepts and an oscillating periodic trend to mimic wet-dry season oscillation. For each ASV, all hosts’ samples were combined into a single series, yielding per-ASV models of CLR dynamics. This procedure is detailed in the Supplemental Methods. Residuals were extracted from these fitted models as seasonally “de-trended” data and CLR correlation matrices across ASV pairs were estimated directly from these adjusted data using cov() in R (**Fig. S9**).

“Synchrony” was estimated by sampling aligned microbiome compositional profiles across hosts. We identified all samples collected from pairs of hosts within 1 calendar day. For instance, a sample collected from host F01 on 2011-03-14 could pair with a sample from M04 on 2011-03-15. For all possible pairs of hosts, we selected one such aligned pair of samples, yielding 1540 joint observations of gut microbiome composition. For each such paired sample, one host was arbitrarily designated as host A and the other as host B. The “synchrony” of a given taxon was estimated as the correlation of a taxon’s model-inferred log-ratio abundance across the set of samples from hosts labeled A and the set of samples from hosts labeled B. The cartoon in **Fig. S10** illustrates this sample pairing.

### Enrichment analyses

We performed enrichment analyses for bacterial families and family pairs in several settings. In each case we computed the frequency of ASVs belonging to a given family, or of pairs belonging to a family pair, on a subset of the data. These were compared to the overall frequencies of ASVs belonging to those families or pairs.

To determine the enrichment of families and family pairs in the most universal ASV pairs (**Fig. 3E and 3F**), we calculated the frequencies of ASV families and pairs in the top 2.5% of pairs by universality scores. Significant enrichment of families or pairs was identified using a one-sided Fisher’s Exact Test. Multiple test correction was applied as a Benjamini-Hochberg adjustment to observed p-values.

Phylogenetic distances between ASV sequences were calculated with the dist.ml function in the “phangorn” package in R [61] using default settings for amino acid substitution rates. In **Fig. 4C and 4D**, low phylogenetic distance/high median correlation strength pairs were identified as those with phylogenetic distances of less than 0.2 and median correlation strengths of greater than 0.5. Again, significance of these was evaluated against overall frequencies of the same families and pairs.

To determine enrichment of low synchrony/high universality or of high synchrony/high universality families and pairs (shown in **Fig. S12A and 12B**), we defined the low synchrony/high universality cohort as those ASV pairs with synchrony estimates of less than 0.3 and universality estimates greater than 0.4. We defined the high synchrony/high universality cohort as those ASV pairs with synchrony greater than 0.3 and universality greater than 0.4. The frequency of these subsets was evaluated against the overall frequencies of the same families and pairs.

### Evaluating explanatory factors

#### Variation in taxon-taxon correlation patterns explained by kinship and baseline composition

To evaluate a possible explanatory effect of distances in terms of kinship or baseline gut bacterial composition on distances in terms of taxon-taxon correlation patterns, we applied Mantel tests. However, because population structure can lead to anticonservative p-values [62], we also developed a second simulation-based procedure for evaluating the significance of baseline composition, using a permutation procedure of our own design. Firstly, baseline composition for each host was estimated by transforming all of a given host’s samples to the centered log-ratio representation after adding a small fraction (0.5) to remove zeros. The vector of per-taxon averages of these CLR values was used as that host’s “baseline” CLR composition. The Euclidean distances between hosts in terms of these per-host baselines were compared against distances in terms of correlation patterns to give an r^2^ value.

In the case of the customized permutation test, this observed result was evaluated against a pseudo-null distribution computed in the following way. The identity of each taxon in the baseline composition was shuffled for each host independently. Euclidean distances across these shuffled baselines were computed and an r^2^ value calculated for these distances against the observed distances computed from taxon-taxon correlation patterns. This procedure was repeated 1000 times to give a distribution of “random” r^2^ values we used as an empirical null.

#### Variation in taxon-taxon correlation patterns explained by sex and social group

To test whether host sex or social group membership predicted similarity in terms of correlation patterns, we used an ANOVA-like strategy. We calculated the F-statistic, a ratio of between-to within-group variation, on the observed correlation patterns (strictly, the vectorized CLR taxon-taxon correlation matrices; *Z* in the equation below) and segmented samples into groups defined by either sex or social group. The F-statistic was calculated as

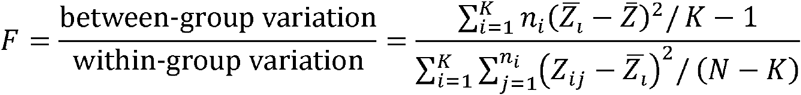

and significance was evaluated via an F-distribution parameterized by the appropriate degrees of freedom. Here *K* represents the number of groups (e.g. two, in the case of sex) and *N*, the total number of hosts. The matrix 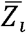 consists of the mean taxon-taxon correlations for group *i* and 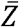, the population mean correlations.

### Comparison to microbiome time series from human populations

We compared our findings to those generated from two human data sets: the DIABIMMUNE project’s infant/toddler cohort from Russian Karelia [44] and the adult diet-microbiome association study of Johnson et al. [34]. In both cases, count tables were obtained from the project’s public website and subject identity and sampling schedules were available in the associated metadata. We compared each host cohort’s universality at the family/order/class level because this taxonomic level offered the greatest comparative power (10.1% of families/orders/classes overlap between the cohorts compared to just 3.1% of genera and no ASVs). The *basset* model from the “fido” R package [46] was fit to each subject’s data set using model settings analogous to those employed on the Amboseli baboon series: first, only taxa with non-zero counts in at least 20% of all subjects’ series were retained; second, Gaussian process kernel bandwidth settings were chosen in such a way as to encode an expectation of minimum autocorrelation between samples at a distance in time of 90 days. We extracted centered log-ratio estimates of taxa at the family level in the same manner as described previously for the Amboseli data set.

## Acknowledgements

We thank Jeanne Altmann for her essential role in stewarding the Amboseli Baboon Project, and in collecting and maintaining the fecal samples used in this manuscript. We thank the Kenya Wildlife Service, the National Council for Science, Technology, and Innovation, and the National Environment Management Authority for permission to conduct research and collect biological samples in Kenya. We also thank the University of Nairobi, Institute of Primate Research, National Museums of Kenya, the Amboseli-Longido pastoralist communities, the Enduimet Wildlife Management Area, Ker & Downey Safaris, Air Kenya, and Safarilink for their cooperation and assistance in the field. We thank Karl Pinc for managing and designing the database. We also thank Tawni Voyles, Anne Dumaine, Yingying Zhang, Meghana Rao, Tauras Vilgalys, Amanda Lea, Noah Snyder-Mackler, Paul Durst, Jay Zussman, Garrett Chavez, and Reena Debray for contributing to fecal sample processing. Complete acknowledgments for the ABRP can be found online at https://amboselibaboons.nd.edu/acknowledgements/.

## Funding

This work was supported by the National Science Foundation and the National Institutes of Health, especially NSF Rules of Life Award DEB 1840223 (EAA, JAG), the National Institute on Aging for R01 AG071684 (EAA), R21 AG055777 (EAA, RB), NIH R01 AG053330 (EAA), and NIH R35 GM128716 (RB). We also thank the Duke University Population Research Institute P2C-HD065563 (pilot to JT), the University of Notre Dame’s Eck Institute for Global Health (EAA), and the Notre Dame Environmental Change Initiative (EAA). Since 2000, long-term data collection in Amboseli has been supported by NSF and NIH, including IOS 1456832 (SCA), IOS 1053461 (EAA), DEB 1405308 (JT), IOS 0919200 (SCA), DEB 0846286 (SCA), DEB 0846532 (SCA), IBN 0322781 (SCA), IBN 0322613 (SCA), BCS 0323553 (SCA), BCS 0323596 (SCA), P01AG031719 (SCA), R21AG049936 (JT, SCA), R03AG045459 (JT, SCA), R01AG034513 (SCA), R01HD088558 (JT), and P30AG024361 (SCA). We also thank Duke University, Princeton University, the University of Notre Dame, the Chicago Zoological Society, the Max Planck Institute for Demographic Research, the L.S.B. Leakey Foundation and the National Geographic Society for support at various times over the years.

## Data and code availability

16S rRNA gene sequences are available on EBI-ENA (project 590 ERP119849) and Qiita (study 12949). Analyzed data and code is available on GitHub at: https://github.com/kimberlyroche/rulesoflife

## Notes

### Competing Interest Statement

The authors have declared no competing interest.

### Summary of Updates

Acknowledgements section added

## References

1. Faust K, Raes J. Microbial interactions: from networks to models. Nature Reviews Microbiology. 2012;10(8):538–50. doi: 10.1038/nrmicro2832. PubMed PMID: 22796884.

2. Foster KR, Bell T. Competition, Not Cooperation, Dominates Interactions among Culturable Microbial Species. Current Biology. 2012;22(19):1845–50. doi: 10.1016/j.cub.2012.08.005. PubMed PMID: WOS:000309792900031.

3. Dolinsek J, Goldschmidt F, Johnson DR. Synthetic microbial ecology and the dynamic interplay between microbial genotypes. Fems Microbiology Reviews. 2016;40(6):961–79. doi: 10.1093/femsre/fuw024. PubMed PMID: WOS:000387995000010.

4. Seth EC, Taga ME. Nutrient cross-feeding in the microbial world. Front Microbiol. 2014;5:350. Epub 20140708. doi: 10.3389/fmicb.2014.00350. PubMed PMID: 25071756; PubMed Central PMCID: PMCPMC4086397.

5. Backhed F, Ley RE, Sonnenburg JL, Peterson DA, Gordon JI. Host-bacterial mutualism in the human intestine. Science. 2005;307(5717):1915–20. doi: Doi 10.1126/Science.1104816. PubMed PMID: ISI:000227957300045.

6. Gould AL, Zhang V, Lamberti L, Jones EW, Obadia B, Korasidis N, et al. Microbiome interactions shape host fitness. Proc Natl Acad Sci U S A. 2018. doi: 10.1073/pnas.1809349115. PubMed PMID: 30510004.

7. Pontrelli S, Szabo R, Pollak S, Schwartzman J, Ledezma-Tejeida D, Cordero OX, et al. Metabolic cross-feeding structures the assembly of polysaccharide degrading communities. Sci Adv. 2022;8(8):eabk3076. Epub 20220223. doi: 10.1126/sciadv.abk3076. PubMed PMID: 35196097; PubMed Central PMCID: PMCPMC8865766.

8. Degnan PH, Taga ME, Goodman AL. Vitamin B12 as a modulator of gut microbial ecology. Cell Metab. 2014;20(5):769–78. doi: 10.1016/j.cmet.2014.10.002. PubMed PMID: 25440056; PubMed Central PMCID: PMCPMC4260394.

9. Loftus M, Hassouneh SA, Yooseph S. Bacterial associations in the healthy human gut microbiome across populations. Sci Rep. 2021;11(1):2828. Epub 20210202. doi: 10.1038/s41598-021-82449-0. PubMed PMID: 33531651; PubMed Central PMCID: PMCPMC7854710.

10. Bashan A, Gibson TE, Friedman J, Carey VJ, Weiss ST, Hohmann EL, et al. Universality of human microbial dynamics. Nature. 2016;534(7606):259–62. doi: 10.1038/nature18301. PubMed PMID: 27279224; PubMed Central PMCID: PMCPMC4902290.

11. Widder S, Allen RJ, Pfeiffer T, Curtis TP, Wiuf C, Sloan WT, et al. Challenges in microbial ecology: building predictive understanding of community function and dynamics. ISME Journal. 2016;10(11):2557–68. doi: 10.1038/ismej.2016.45. PubMed PMID: 27022995.

12. Cao HT, Gibson TE, Bashan A, Liu YY. Inferring human microbial dynamics from temporal metagenomics data: Pitfalls and lessons. Bioessays. 2017;39(2):1600188. doi: 10.1002/bies.201600188. PubMed PMID: 28000336.

13. Faust K, Raes J. Host-microbe interaction: Rules of the game for microbiota. Nature. 2016;534(7606):182–3. doi: 10.1038/534182a. PubMed PMID: 27279206.

14. Coyte KZ, Schluter J, Foster KR. The ecology of the microbiome: Networks, competition, and stability. Science. 2015;350(6261):663–6. doi: 10.1126/science.aad2602. PubMed PMID: 26542567.

15. Palmer JD, Foster KR. Bacterial species rarely work together. Science. 2022;376(6593):581–2. Epub 20220505. doi: 10.1126/science.abn5093. PubMed PMID: 35511986.

16. Wu G, Zhao N, Zhang C, Lam YY, Zhao L. Guild-based analysis for understanding gut microbiome in human health and diseases. Genome Med. 2021;13(1):22. Epub 2021/02/11. doi: 10.1186/s13073-021-00840-y. PubMed PMID: 33563315; PubMed Central PMCID: PMCPMC7874449.

17. Coyte KZ, Rao C, Rakoff-Nahoum S, Foster KR. Ecological rules for the assembly of microbiome communities. PLoS Biol. 2021;19(2):e3001116. Epub 20210219. doi: 10.1371/journal.pbio.3001116. PubMed PMID: 33606675; PubMed Central PMCID: PMCPMC7946185.

18. Gao C, Montoya L, Xu L, Madera M, Hollingsworth J, Purdom E, et al. Fungal community assembly in drought-stressed sorghum shows stochasticity, selection, and universal ecological dynamics. Nat Commun. 2020;11(1):34. Epub 2020/01/09. doi: 10.1038/s41467-019-13913-9. PubMed PMID: 31911594; PubMed Central PMCID: PMCPMC6946711.

19. Vila JCC, Liu YY, Sanchez A. Dissimilarity-Overlap analysis of replicate enrichment communities. ISME J. 2020;14(10):2505–13. Epub 20200618. doi: 10.1038/s41396-020-0702-7. PubMed PMID: 32555503; PubMed Central PMCID: PMCPMC7490387.

20. San-Juan-Vergara H, Zurek E, Ajami NJ, Mogollon C, Pena M, Portnoy I, et al. A Lachnospiraceae-dominated bacterial signature in the fecal microbiota of HIV-infected individuals from Colombia, South America. Sci Rep. 2018;8(1):4479. Epub 20180314. doi: 10.1038/s41598-018-22629-7. PubMed PMID: 29540734; PubMed Central PMCID: PMCPMC5852036.

21. Gonze D, Coyte KZ, Lahti L, Faust K. Microbial communities as dynamical systems. Current Opinion in Microbiology. 2018;44:41–9. doi: 10.1016/j.mib.2018.07.004. PubMed PMID: WOS:000447581000007.

22. Franzosa EA, Huang K, Meadow JF, Gevers D, Lemon KP, Bohannan BJM, et al. Identifying personal microbiomes using metagenomic codes. Proceedings of the National Academy of Sciences. 2015;112(22):E2930–E8. doi: 10.1073/pnas.1423854112. PubMed PMID: WOS:000355832200014.

23. Faith JJ, Guruge JL, Charbonneau M, Subramanian S, Seedorf H, Goodman AL, et al. The long-term stability of the human gut microbiota. Science. 2013;341(6141):1237439. Epub 2013/07/06. doi: 10.1126/science.1237439. PubMed PMID: 23828941; PubMed Central PMCID: PMC3791589.

24. Bik EM, Costello EK, Switzer AD, Callahan BJ, Holmes SP, Wells RS, et al. Marine mammals harbor unique microbiotas shaped by and yet distinct from the sea. Nat Commun. 2016;7:10516. Epub 20160203. doi: 10.1038/ncomms10516. PubMed PMID: 26839246; PubMed Central PMCID: PMCPMC4742810.

25. Caporaso JG, Lauber CL, Costello EK, Berg-Lyons D, Gonzalez A, Stombaugh J, et al. Moving pictures of the human microbiome. Genome Biology. 2011;12(5):R50. doi: Artn R50 Doi 10.1186/Gb-2011-12-5-R50. PubMed PMID: ISI:000295732700014.

26. Costello EK, Lauber CL, Hamady M, Fierer N, Gordon JI, Knight R. Bacterial community variation in human body habitats across space and time. Science. 2009;326(5960):1694–7. doi: Doi 10.1126/Science.1177486. PubMed PMID: ISI:000272839000053.

27. Louca S, Polz MF, Mazel F, Albright MBN, Huber JA, O’Connor MI, et al. Function and functional redundancy in microbial systems. Nat Ecol Evol. 2018;2(6):936–43. Epub 2018/04/18. doi: 10.1038/s41559-018-0519-1. PubMed PMID: 29662222.

28. Rainey PB, Quistad SD. Toward a dynamical understanding of microbial communities. Philos Trans R Soc Lond B Biol Sci. 2020;375(1798):20190248. Epub 2020/03/24. doi: 10.1098/rstb.2019.0248. PubMed PMID: 32200735; PubMed Central PMCID: PMCPMC7133524.

29. Martiny JB, Jones SE, Lennon JT, Martiny AC. Microbiomes in light of traits: A phylogenetic perspective. Science. 2015;350(6261):aac9323. doi: 10.1126/science.aac9323. PubMed PMID: 26542581.

30. Debray R, Herbert RA, Jaffe AL, Crits-Christoph A, Power ME, Koskella B. Priority effects in microbiome assembly. Nat Rev Microbiol. 2022;20(2):109–21. Epub 20210827. doi: 10.1038/s41579-021-00604-w. PubMed PMID: 34453137.

31. Risely A, Wilhelm K, Clutton-Brock T, Manser MB, Sommer S. Diurnal oscillations in gut bacterial load and composition eclipse seasonal and lifetime dynamics in wild meerkats. Nat Commun. 2021;12(1):6017. Epub 20211014. doi: 10.1038/s41467-021-26298-5. PubMed PMID: 34650048; PubMed Central PMCID: PMCPMC8516918.

32. Kolodny O, Weinberg M, Reshef L, Harten L, Hefetz A, Gophna U, et al. Coordinated change at the colony level in fruit bat fur microbiomes through time. Nature Ecology & Evolution. 2019;3(1): 116–24. doi: 10.1038/s41559-018-0731-z. PubMed PMID: WOS:000453767000021.

33. Flores GE, Caporaso JG, Henley JB, Rideout JR, Domogala D, Chase J, et al. Temporal variability is a personalized feature of the human microbiome. Genome Biology. 2014;15(12):531. doi: ARTN 531 10.1186/s13059-014-0531-y. PubMed PMID: WOS:000346609500011.

34. Johnson AJ, Vangay P, Al-Ghalith GA, Hillmann BM, Ward TL, Shields-Cutler RR, et al. Daily Sampling Reveals Personalized Diet-Microbiome Associations in Humans. Cell Host & Microbe. 2019;25(6):789–802. Epub 2019/06/14. doi: 10.1016/j.chom.2019.05.005. PubMed PMID: 31194939.

35. Smits SA, Marcobal A, Higginbottom S, Sonnenburg JL, Kashyap PC. Individualized Responses of Gut Microbiota to Dietary Intervention Modeled in Humanized Mice. mSystems. 2016.

36. Kalyuzhny M, Shnerb NM. Dissimilarity-overlap analysis of community dynamics: Opportunities and pitfalls. Methods in Ecology and Evolution. 2017;8:1764–73.

37. Marsland R, 3rd, Cui W, Mehta P. A minimal model for microbial biodiversity can reproduce experimentally observed ecological patterns. Sci Rep. 2020;10(1):3308. Epub 20200224. doi: 10.1038/s41598-020-60130-2. PubMed PMID: 32094388; PubMed Central PMCID: PMCPMC7039880.

38. Faust K, Lahti L, Gonze D, de Vos WM, Raes J. Metagenomics meets time series analysis: unraveling microbial community dynamics. Current Opinion in Microbiology. 2015;25:56–66. doi: 10.1016/j.mib.2015.04.004. PubMed PMID: 26005845.

39. Gloor GB, Macklaim JM, Pawlowsky-Glahn V, Egozcue JJ. Microbiome datasets are compositional: and this is not optional. Front Microbiol. 2017;8:2224. Epub 2017/12/01. doi: 10.3389/fmicb.2017.02224. PubMed PMID: 29187837; PubMed Central PMCID: PMCPMC5695134.

40. Quinn TP, Richarrson MF, Lovell D, Crowley TM. propr: An R-package for Identifying Proportionally Abundant Features Using Compositional Data Analysis Scientific Reports. 2017;7:16252.

41. Alberts SC, Altmann J. The Amboseli Baboon Research Project: Themes of continuity and change. In: Kappeler P, Watts DP, editors. Long-term field studies of primates: Springer Verlag; 2012. p. 26187.

42. Bjo□rk J, Dasari M, Grieneisen L, Gould TJ, Grenier JC, Yotova V, et al. Synchrony and idiosyncrasy in the gut microbiome of wild primates. Nature Ecology & Evolution. 2022;6:955–64. doi: https://www.biorxiv.org/content/10.1101/2021.11.24.469913v1.

43. Grieneisen L, Dasari M, Gould TJ, Bjo□rk JR, Grenier JC, Yotova V, et al. Gut microbiome heritability is near-universal but environmentally contingent. Science. 2021;373:181–6.

44. Vatanen T, Kostic AD, d’Hennezel E, Siljander H, Franzosa EA, Yassour M, et al. Variation in Microbiome LPS Immunogenicity Contributes to Autoimmunity in Humans. Cell. 2016;165(4):842–53. Epub 20160428. doi: 10.1016/j.cell.2016.04.007. PubMed PMID: 27133167; PubMed Central PMCID: PMCPMC4950857.

45. Cullen CM, Aneja KK, Beyhan S, Cho CE, Woloszynek S, Convertino M, et al. Emerging Priorities for Microbiome Research. Front Microbiol. 2020;11:136. Epub 2020/03/07. doi: 10.3389/fmicb.2020.00136. PubMed PMID: 32140140; PubMed Central PMCID: PMCPMC7042322.

46. Silverman JD, Roche K, Holmes ZC, David LA, Mukherjee S. Bayesian Multinomial Logistic Normal Models through Marginally Latent Matrix-T Processes. Journal of Machine Learning Research. 2022;23:1–42.

47. Vetrovsky T, Baldrian P. The variability of the 16S rRNA gene in bacterial genomes and its consequences for bacterial community analyses. PLoS One. 2013;8(2):e57923. Epub 20130227. doi: 10.1371/journal.pone.0057923. PubMed PMID: 23460914; PubMed Central PMCID: PMCPMC3583900.

48. Guittar J, Shade A, Litchman E. Trait-based community assembly and succession of the infant gut microbiome. Nat Commun. 2019;10(1):512. Epub 2019/02/03. doi: 10.1038/s41467-019-08377-w. PubMed PMID: 30710083; PubMed Central PMCID: PMCPMC6358638.

49. Kehe J, Ortiz A, Kulesa A, Gore J, Blainey PC, Friedman J. Positive interactions are common among culturable bacteria. Science Advances. 2021;45:eabi7159.

50. Weiss AS, Burrichter AG, Durai Raj AC, von Strempel A, Meng C, Kleigrewe K, et al. In vitro interaction network of a synthetic gut bacterial community. ISME J. 2022;16(4):1095–109. Epub 20211202. doi: 10.1038/s41396-021-01153-z. PubMed PMID: 34857933; PubMed Central PMCID: PMCPMC8941000.

51. Ortiz A, Vega NM, Ratzke C, Gore J. Interspecies bacterial competition regulates community assembly in the C. elegans intestine. ISME J. 2021;15(7):2131–45. Epub 20210215. doi: 10.1038/s41396-021-00910-4. PubMed PMID: 33589765; PubMed Central PMCID: PMCPMC8245486.

52. Carlstrom CI, Field CM, Bortfeld-Miller M, Muller B, Sunagawa S, Vorholt JA. Synthetic microbiota reveal priority effects and keystone strains in the Arabidopsis phyllosphere. Nat Ecol Evol. 2019;3(10):1445–54. Epub 20190926. doi: 10.1038/s41559-019-0994-z. PubMed PMID: 31558832; PubMed Central PMCID: PMCPMC6774761.

53. Venturelli OS, Carr AC, Fisher G, Hsu RH, Lau R, Bowen BP, et al. Deciphering microbial interactions in synthetic human gut microbiome communities. Mol Syst Biol. 2018;14(6):e8157. Epub 20180621. doi: 10.15252/msb.20178157. PubMed PMID: 29930200; PubMed Central PMCID: PMCPMC6011841.

54. Meehan CJ, Beiko RG. A phylogenomic view of ecological specialization in the Lachnospiraceae, a family of digestive tract-associated bacteria. Genome Biol Evol. 2014;6(3):703–13. doi: 10.1093/gbe/evu050. PubMed PMID: 24625961; PubMed Central PMCID: PMCPMC3971600.

55. Vacca M, Celano G, Calabrese FM, Portincasa P, Gobbetti M, De Angelis M. The Controversial Role of Human Gut Lachnospiraceae. Microorganisms. 2020;8(4). Epub 20200415. doi: 10.3390/microorganisms8040573. PubMed PMID: 32326636; PubMed Central PMCID: PMCPMC7232163.

56. Stackebrandt E. The Family Lachnospiraceae. The Prokaryotes. 2014.

57. Costello EK, Stagaman K, Dethlefsen L, Bohannan BJ, Relman DA. The application of ecological theory toward an understanding of the human microbiome. Science. 2012;336(6086):1255–62. Epub 2012/06/08. doi: science.1224203 [pii] 10.1126/science.1224203. PubMed PMID: 22674335.

58. Walter J, Ley R. The human gut microbiome: ecology and recent evolutionary changes. Annual Review of Microbiology. 2011;65:411–29. Epub 2011/06/21. doi: 10.1146/annurev-micro-090110-102830. PubMed PMID: 21682646.

59. McMurdie PJ, Holmes S. phyloseq: an R package for reproducible interactive analysis and graphics of microbiome census data. PLoS One. 2013;8(4):e61217. Epub 20130422. doi: 10.1371/journal.pone.0061217. PubMed PMID: 23630581; PubMed Central PMCID: PMCPMC3632530.

60. Yaqing Chen [aut c, Alvaro Gajardo [aut], Jianing Fan [aut], Zhong Q, Dubey P, Han K, et al. frechet: Statistical Analysis for Random Objects and Non-Euclidean Data. Available: https://CRANR-projectorg/package=frechet. 2020.

61. Schliep KP. phangorn: phylogenetic analysis in R. Bioinformatics. 2011;27(4):592–3. Epub 20101217. doi: 10.1093/bioinformatics/btq706. PubMed PMID: 21169378; PubMed Central PMCID: PMCPMC3035803.

62. Guillot G, Rousset F. Dismantling the Mantel tests. In: Harmon L, editor. Methods in Ecology and Evolution. 42013. p. 336–44.

