## Supplemental Information for "Universal gut microbial relationships in the gut microbiome of wild baboons"

Supplemental materials for ***Universal gut microbial relationships in the gut microbiome of wild baboons***

### Supplemental Methods

The majority of our methodological details are in the Methods section of the main text. Below we provide more information on our procedures for de-trending taxa for potential seasonal effects and for estimating synchronized dynamics between taxa.

#### De-trending for season

Seasonally de-trended data was obtained in the following way. The observed ASV count matrices were centered log-ratio transformed and linear autoregressive models were fit to each centered log-transformed (CLR) amplicon sequence variant’s (ASV) series. In these models, wet-dry season oscillation was modeled as a sine wave with a period of 365 days. The magnitude of this component was estimated during model fitting after an offset (in weeks) was estimated in a first step, in order to best align the oscillating seasonal component with the data. Per-ASV models were fit using the following syntax:

arima(x = x, xreg = model.matrix(~ factor(host) + sin_f(offset, days))[,-1], order = c(1, 0, 0))

where x are CLR counts for that ASV, the “order” argument of arima enables a single autoregressive component, and the “xreg” argument specifies a covariance matrix. That covariance matrix contains a per-host label giving host-specific offsets for log-ratio abundance and an oscillating seasonal trend through “sin_f,” a function that samples values corresponding to day indices (through “days”) from a sine wave with weekly offset (“offset”) and a period of 365 days. The residuals from these per-ASV model fits were extracted and used as the seasonally “de-trended” data (see **Fig. S10B**).

Correlations across CLR ASV-ASV pairs were estimated from these residual series with the cov() function in R.

#### Estimating synchrony

The high universality scores we observed for some ASV pairs in Amboseli could arise if the individual members of an ASV-ASV pair are strongly synchronized over time in different hosts, even if the pair does not directly interact. That is, if one member of the pair is highly synchronized with itself in different hosts, and the other member is also strongly synchronized with itself, then universality could be an inevitable outcome of each member’s strong synchrony.

To test this possibility, we estimated the degree to which each of the 125 ASVs (not including rare ASVs) exhibited synchronized dynamics across the 56 hosts by first identifying pairs of samples from different hosts collected within 24 hours of each other and randomly selecting a single same-day pair per host pair. This procedure produced 1540 same-day sample pairs across hosts. One partner in these pairs was arbitrarily assigned to series “A” and the other, to series “B” (**Fig. S10A**). The correlation of model-inferred log-ratio abundance across series A and B was calculated to give an estimate of the synchrony of log-ratio change in this ASV across the population and across time (**Fig. S10B**) as an estimate of the degree of shared perturbation (**Fig. S10C**). The observed estimates are entirely positive. Repeating this procedure where paired samples have had their order shuffled in time independently across hosts results in near-zero estimates of synchrony (**Figs. S10D**).

### Supplemental Results

#### Universality in Amboseli is only weakly explained by synchrony

As discussed above in “Estimating synchrony”, universality could be an inevitable outcome of each member’s strong synchrony. We estimated synchronous ASV-level change across the population by using the procedure described above. To derive expectations of “null” synchrony, we repeated the same procedure on data where sample order was shuffled (**Fig. S10**).

We found that 94.4% of ASVs (118 of 125) had synchrony estimates that were higher than expected (FDR ≤ 0.05, permutation testing; **Fig. S10D**; **Table S8**). However, consistent with prior research on this data set [1], synchrony estimates were still modest, with median synchrony across all ASVs of 0.187 (range = 0.00483 to 0.480; **Fig. S10D**; **Table S8**). The most synchronized ASV (#21) belonged to the family Clostridiaceae 1; its synchrony (r = 0.480) across five especially well-sampled hosts is visualized in **Fig. S10C**. Only six other ASVs (<5%; 6 of 125) exhibited synchrony >0.4, including ASV #22, in genus Oribacterium, ASVs #106 and #107 in family Lachnospiraceae, ASV #50 in the family Bifidobacteriaceae, and ASV #115 in family Coriobacteriaceae. Interestingly, all these most synchronized ASVs are included in the large cluster in the network of most universal ASV pairs in **Fig. 3D**, and ASV #107 is also the most connected ASV in this network. The five taxa with the lowest synchrony estimates included a member genera Libanicoccus (ASV#17) and Brachyspira (ASV #102), two members of the order Clostridiales (ASVs #39 and #91), and a member of order Bacteriodales (ASV #98; **Tables S1 and S8**).

To understand the degree to which the universality scores for ASV-pairs in our data set could be attributed to synchronized dynamics at the level of individual members of the pair, we correlated the universality score of each pair with the average synchrony for members of the pair (**Fig. S12A**). In support of the idea that universality is not solely driven by synchrony, the Pearson’s correlation for the relationship between universality and mean synchrony was 0.264. Some pairs exhibited especially high universality, despite low mean synchrony, suggesting they may interact directly and do so predictably across hosts (upper left corner of **Fig. S12A**). These pairs were enriched for members of the families Atopobiaceae, Eggerthellaceae, Erysipelotrichaceae, and Lachnospiraceae (Fisher’s Exact Test, p < 0.0001, FDR ≤ 0.05; **Fig. S12B**; **Table S9**). Other pairs exhibited high universality and high mean synchrony (upper right corner of **Fig. S12A**). These pairs were enriched for members of the families Bifidobacteriaceae, Lachnospiraceae, Ruminococcaceae (Fisher’s Exact Test, p < 0.0001; FDR ≤ 0.05) and others; **Fig. S12B**; **Table S9**).

#### Estimates of universality are not strongly affected by taxon relative abundance

We were concerned that differences in average abundances could play a role in estimates of “universality.” In particular, less abundant ASVs may drop out of sampling more often and this competition with more abundant ASVs to be sampled may cause these rare taxa to be more frequently negatively correlated with others.

We correlated the average abundance of ASVs with the median correlation strength observed for pairs in which that ASV was a member and observed a small but significant association between log-ratio abundance and median correlation strength. Increased average CLR abundance of the *more common partner in a taxon pair* was associated with a small increase in the median correlation strength (r = 0.012, p < 0.0001; **Fig. S6A**).

Next, we examined whether the pairwise abundance characteristics of pairs in the top 2.5% by universality score differed from all other pairs. The less abundant partner in pairs in the top 2.5% by universality score had a slightly higher average abundance than did the less partner in the other 97.5% of pairs (Wilcoxon rank sum test, difference=0.153, p=0.00376; **Fig. S6B**).

Differences in mean CLR abundance are slightly smaller in the top 2.5% of pairs than in other pairs (Wilcoxon rank sum test, difference=-0.214, p = 0.0009; **Fig. S6C**).

#### Choice of minimum informative sample number per-host

The full set of sequenced Amboseli baboon 16S rRNA samples consists of 17,265 samples from 600 individual hosts, previously analyzed in Grieneisen et al [2] and Björk et al. [1]. Because many of the 600 hosts in this full data set were sampled only a handful of times, we subset our analysis to focus especially densely sampled individuals. To determine which hosts to target, we began by estimating the minimum number of samples per host required to reliably estimate microbe-microbe correlations. We simulated data from our generative model (see main text Methods) in order to estimate this minimum informative sample number. Sequence counts for 126 synthetic “taxa” were simulated across 1000 continuous “days” of sampling. These taxa covaried according to randomly generated patterns of covariance.

We randomly downsampled the simulated series to include 10, 20, 30, 40, 50, 75, 100, or 200 samples per host, and then fit our model on each of these downsampled data sets, extracting maximum a posteriori (MAP) estimates of the correlation matrix over centered log-ratio taxa. To score the fidelity of the information retrieved by the model, we calculated the percent agreement in sign between the estimated and true (simulated) taxon-taxon covariance patterns.

We found that downsampled simulations that included at least 75 samples retained sufficient information for the model to recover 90% or more of simulated correlations, in terms of the sign of the correlation (**Fig. S13**). We selected this threshold because it provides a reasonable tradeoff between accuracy of estimation and power for our main analyses, resulting in a final analysis data set of 56 hosts and 5,534 samples.

#### Sensitivity of estimates to kernel hyperparameters

Our Gaussian process model uses the following parameterization for its kernels. The *bandwidth* for a squared exponential kernel (which manages sample-sample autocorrelation) is chosen such that this autocorrelation decays to a minimum at 90 days. This mirrors the behavior of empirical estimates of sample-sample autocorrelation in the data. A second kernel models sample-sample covariance driven by gross similarity in diet (as encoded by the top principal components of dietary composition for the population) and constitutes 25% of total sample-sample variation.

We fit four alternative versions of our models in order to test the sensitivity of these parameter settings, varying the bandwidth of the squared exponential kernel in such a way as to give minimum sample-sample autocorrelation at either 30 or 90 days. We also varied the proportion of sample-sample covariance driven by diet from 0% to 25% to 50%, and we varied the log scale of total sample-sample variance between 1 and 2. In all cases, estimates of correlation between CLR ASVs were similar, with minimum and maximum r^2^ between “canonical” and alternative model estimates of 0.993 and 0.996 respectively. This suggests our findings are reasonably robust to a range of hyperparameter settings.

#### Comparison to COAT estimates of dynamics

Because the *basset* model estimates centered log-ratio correlation from (potentially biased) estimates of additive log-ratio covariance, we tested the compatibility of *basset* results with those of an alternative model that directly infers CLR covariance. We compared our results to those obtained from COAT [3] which estimates centered log-ratio correlation through a sparsity-inducing procedure that yields more conservative estimates. While COAT estimates are generally more conservative, they also largely agree with *basset* estimates (R = 0.884), including for taxon pairs with the strongest inferred associations (**Fig. S15**).

### Supplemental Figures

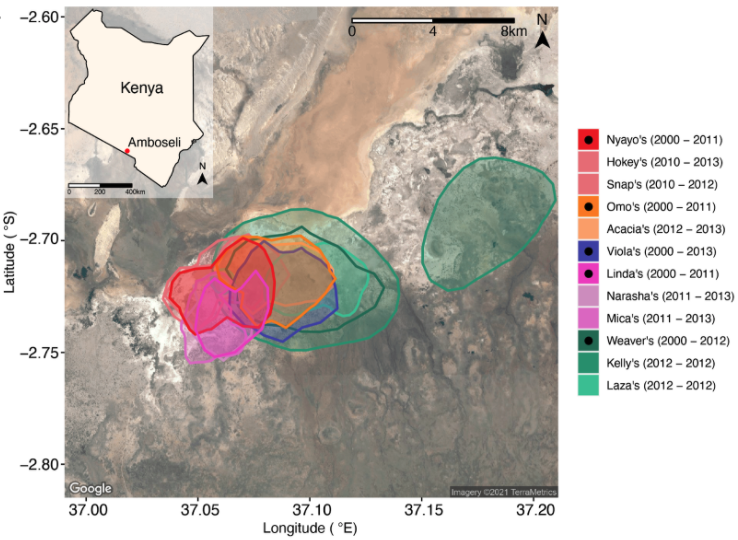

##### Figure S1. Host ranging patterns

Ranging patterns for the baboon social groups included in this study during the period of microbiome sampling (May 19, 2000 to September 19, 2013). See [1] for an in-depth analysis of environmental drivers of microbiome dynamics in this host population.

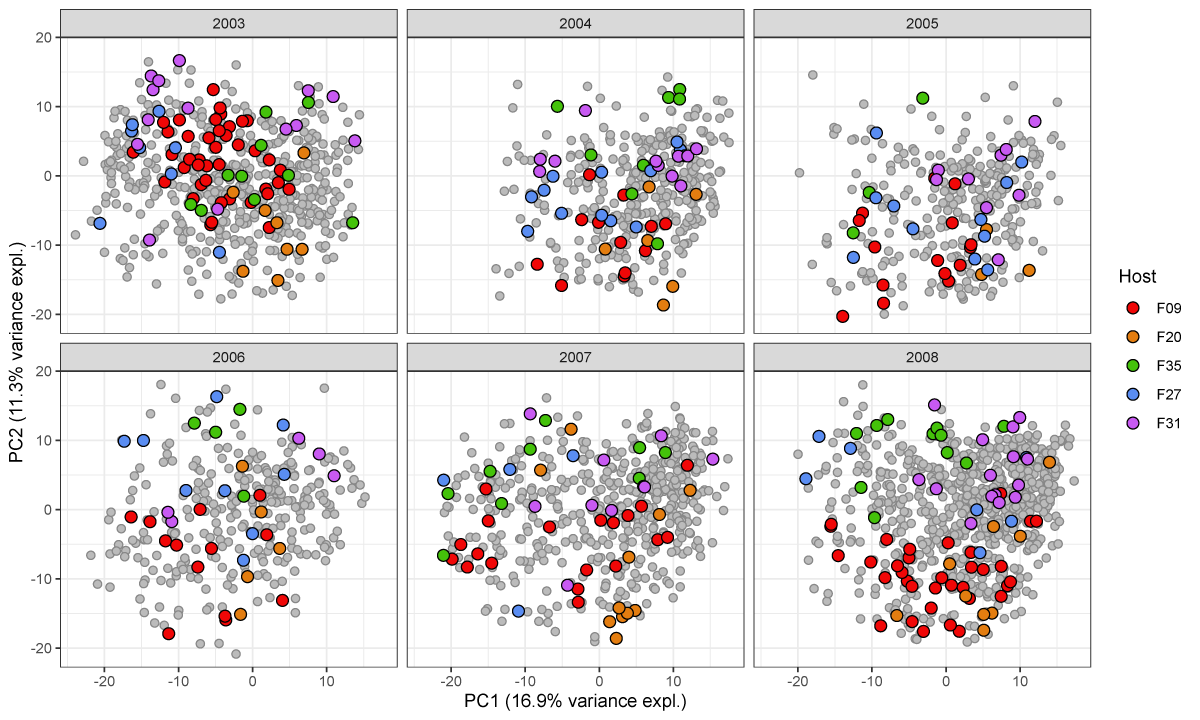

##### Figure S2. Variation across hosts

Each baboon exhibits a somewhat distinctive gut microbial community (p < 0.0001; ANOVA) summarized here by the first two principal components of CLR-transformed ASV abundances of ASVs during six representative years of sample collection (see Fig. 1B). Colored dots represent one of five especially well sampled hosts, and colors indicate samples from the same host. Grey dots represent all other samples collected in the same year.

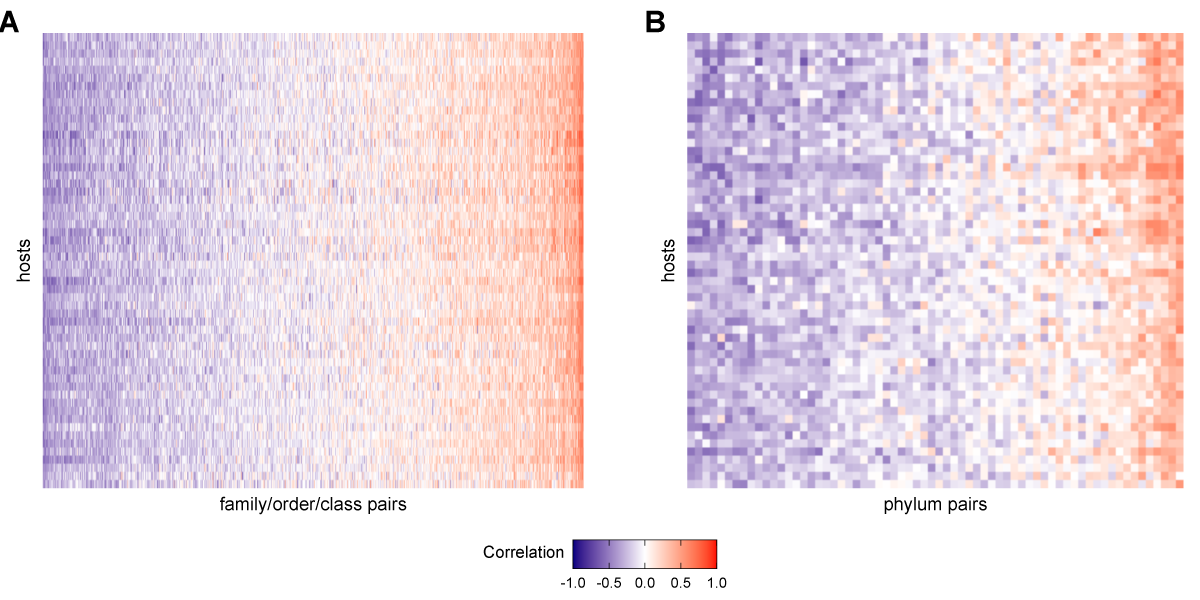

##### Figure S3. Heat maps of correlation of CLR taxon pairs at taxonomic levels higher than ASV

Each heat map shows the Pearson’s correlation coefficient of CLR abundances between all pairs of taxa (x-axis) in each of the 56 baboons (y-axis). Panel **(A)** gives pairs of taxa agglomerated to the most granular possible family, order, or class. Panel **(B)** gives phylum-phylum pairs. See Fig. 2A for a heatmap of ASV-ASV correlation patterns across hosts.

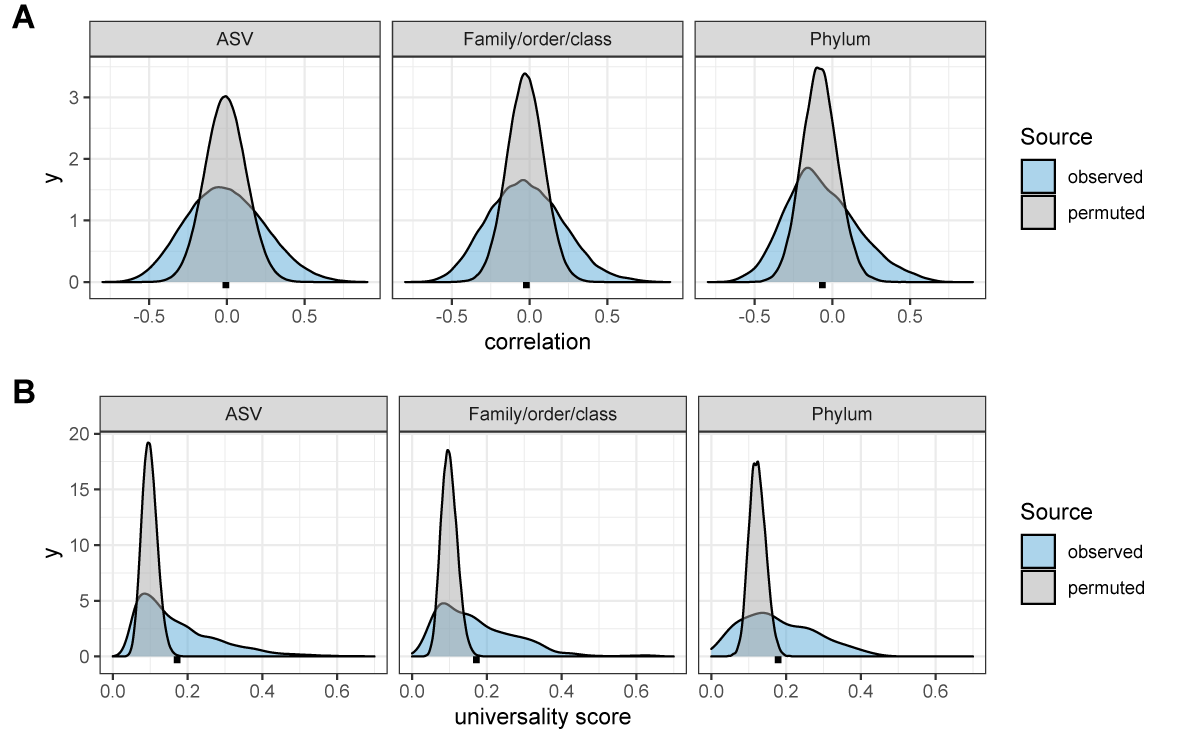

##### Figure S4. Confidence cutoffs for correlations and universality scores

The three plots in panel **(A)** show density distributions of the observed correlations between taxa (blue distributions) compared to random expectation (gray distributions) for three taxonomic partitions of the data: ASVs, family/order/class-level assignments, and phyla. The random distributions were generated by permuting the data 10 times per-host, shuffling taxonomic identities within individual microbiome samples and re-calculating taxon-taxon correlations in these permuted data sets. Mean correlations for the observed distributions are shown as black points below each distribution. The observed correlations for ASVs range from -0.800 to 0.904 (mean = -0.007; median = -0.016); for family/order/class designations range from -0.735 to 0.813 (mean = -0.022; median = -0.031); and for phyla from -0.651 to 0.687 (mean = -0.065; median = -0.092). Panel **(B)** shows density distributions of the observed universality scores in blue compared to random expectations in grey, for pairs of taxa at the ASVs, family/order/class-level, and phyla. Small, dashed black lines underneath each density indicate the mean observed universality score at that taxonomic level.

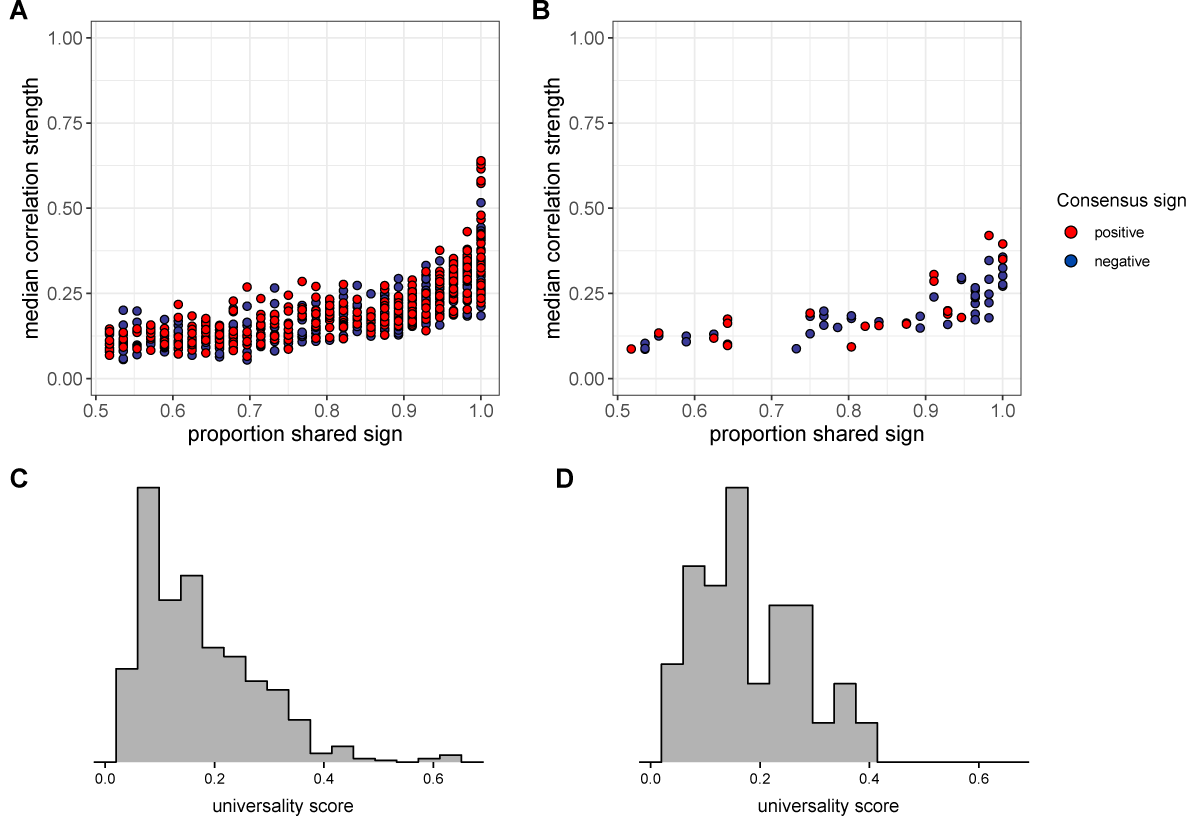

##### Figure S5. Universality at taxonomic levels higher than ASV

The plot in **(A)** shows the median absolute value of each family/order/class pair’s correlation coefficient across hosts as a function of correlation consistency, measured as the proportion of hosts that shared the majority correlation sign (positive or negative; taxon pairs that were positively correlated in half of the 56 hosts have a consistency of 0.5; pairs that were positively [or negatively] correlated in all hosts have a consistency of 1.0). Panel **(B)** plots the same estimates for phylum-phylum pairs. Multiplying the two axes in panels **(A)** and **(B)** creates a universality score reflecting the strength and consistency of pairwise microbial correlations across hosts. Universality score distributions for family/order/class pairs are shown in panel **(C)** and scores for phylum pairs are shown in panel **(D)**.

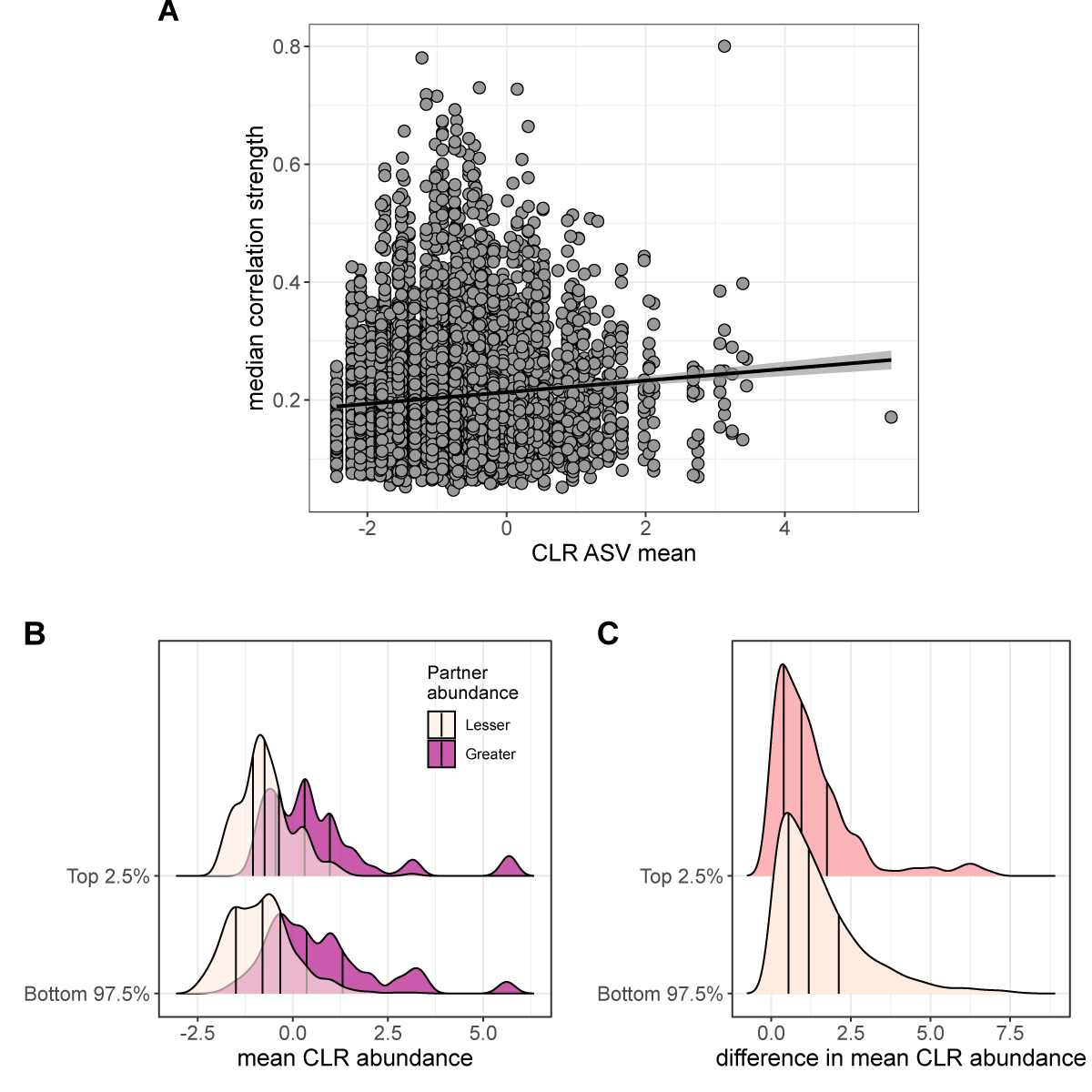

##### Figure S6. Median correlation strength is not strongly linked to bacterial abundance

Panel **(A)**: the mean CLR abundance for the more abundant ASV in each ASV-ASV pair is positively associated with the median strength of correlations across hosts, but this effect is very small (r = 0.012, p-value < 0.0001). The mean CLR abundance of the less abundant partner in the pair was not significantly associated with median correlation strength across hosts (r = -0.0008, p-value = 0.407). Within-pair distributions of mean CLR abundance are also similar in the 2.5% most universal pairs versus all other pairs. Panel **(B)** shows the distribution of observed CLR abundances for pairs of ASVs in the top 2.5% of universality scores compared to the abundances in the remaining 97.5% of pairs. Within each level, light and dark densities show the abundances of the less abundant member of the pair (light) and the more abundant member (dark). The less abundant partner in pairs in the top 2.5% by universality score had a slightly higher average abundance than did the more less partner in the other 97.5% of pairs (Wilcoxon rank sum test, difference=0.153, p=0.00376). Panel **(C)** shows the distributions of pairwise differences in mean CLR abundance for the same subsets of the data in panel **(B)**; the darker pink distribution shows the difference in CLR abundance between ASVs in the top 2.5% most universal pairs compared to the remaining 97.5% of pairs (off-white). Differences are slightly smaller in the top 2.5% of pairs than in other pairs (Wilcoxon rank sum test, difference=-0.214, p = 0.0009).

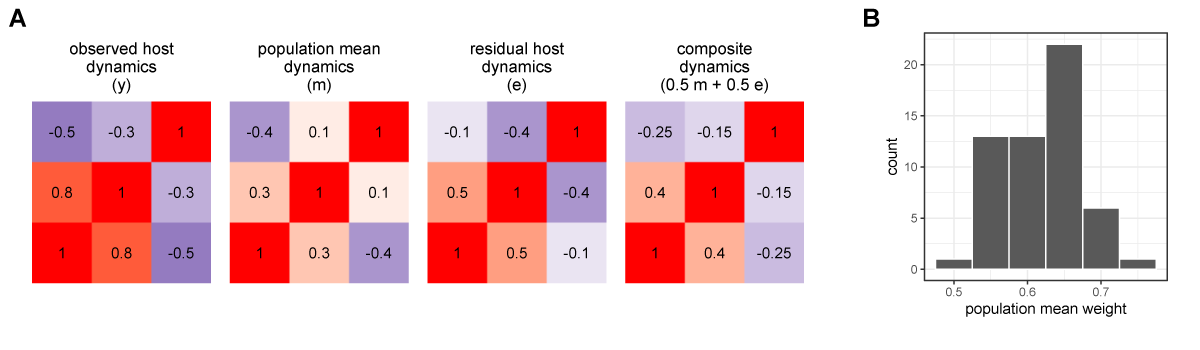

##### Figure S7. Quantifying the relative strength of universal versus individualized dynamics

Panel **(A)** shows observed (y), population mean (m), and host residual dynamics (e = y - m) for a single hypothetical host with three ASVs. The fourth matrix shows a simulated case where host-level and population-level mean dynamics are added in equal proportions (0.5 m + 0.5 e) to approximate y. We estimate the proportion host- versus individual-level signal as the combination of mean and residual which best approximates y. Panel **(B)** shows the results of this procedure applied to the Amboseli data, giving a per-host estimate of population-level contribution to the observed CLR ASV-ASV correlations.

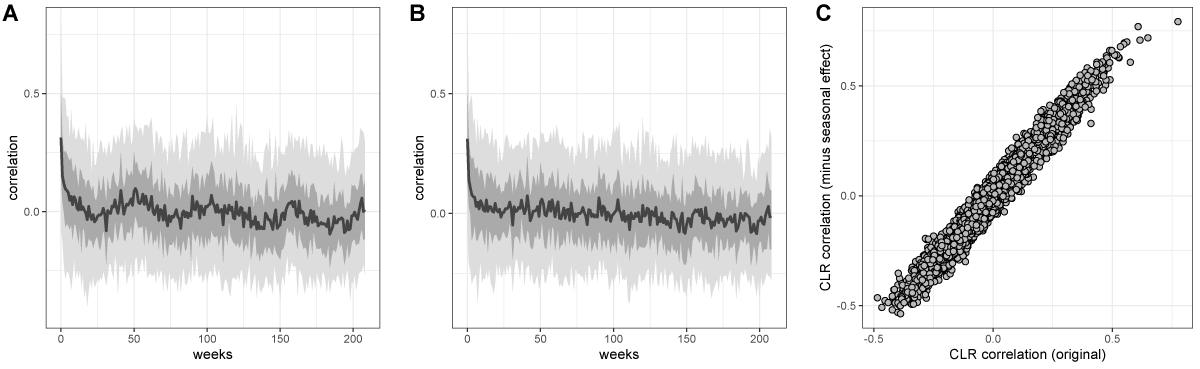

##### Figure S8. Estimates are CLR ASV-ASV correlation are similar after removing seasonal trend

Panel **(A)** shows sample-sample autocorrelation estimates for the CLR-transformed data from 56 baboon hosts. Panel **(B)** shows sample-sample autocorrelation for the same data after the removal of a seasonal trend by arima() in R. Estimates of CLR ASV-ASV correlation derived from the original (basset) model and this per-ASV linear autoregressive model are highly similar, as shown in panel **(C)** where r=0.979.

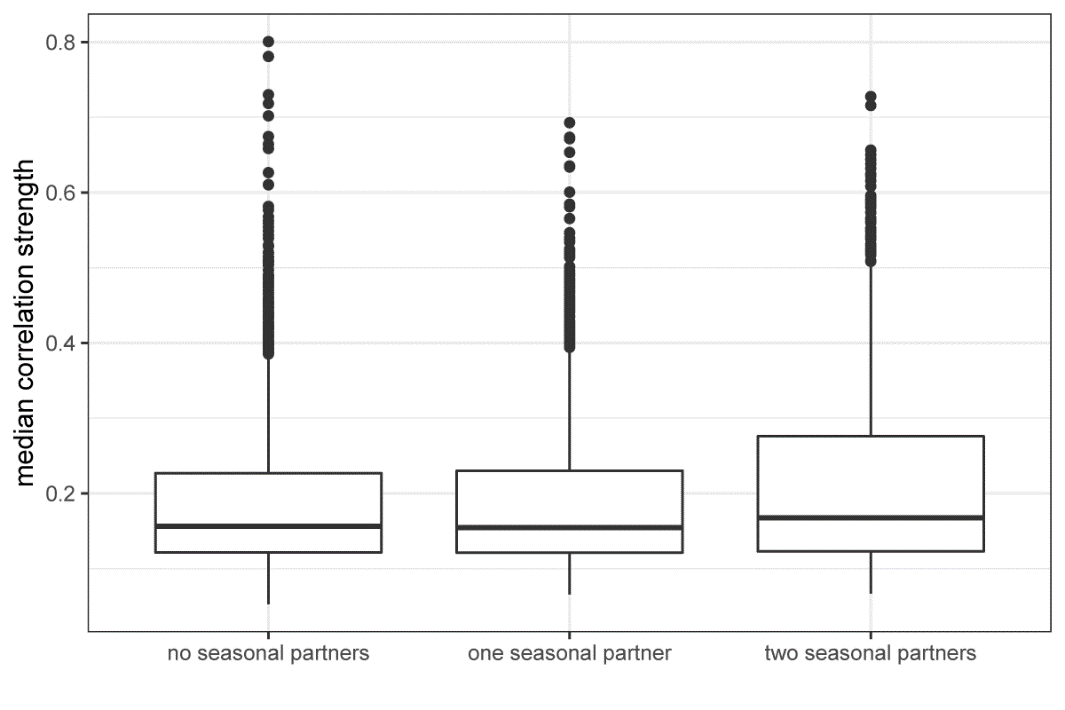

##### Figure S9. ASV-ASV pairs with two “seasonally varying” partners have slightly higher median correlations across hosts

“Seasonally varying” ASVs were those identified as having seasonally differential CLR abundance in Björk et al. (2022). ASV-ASV pairs with two such “seasonal” members had significantly higher median correlation across hosts than did pairs where 1 or 0 partners were seasonal (difference of 0.026, p < 0.0001).

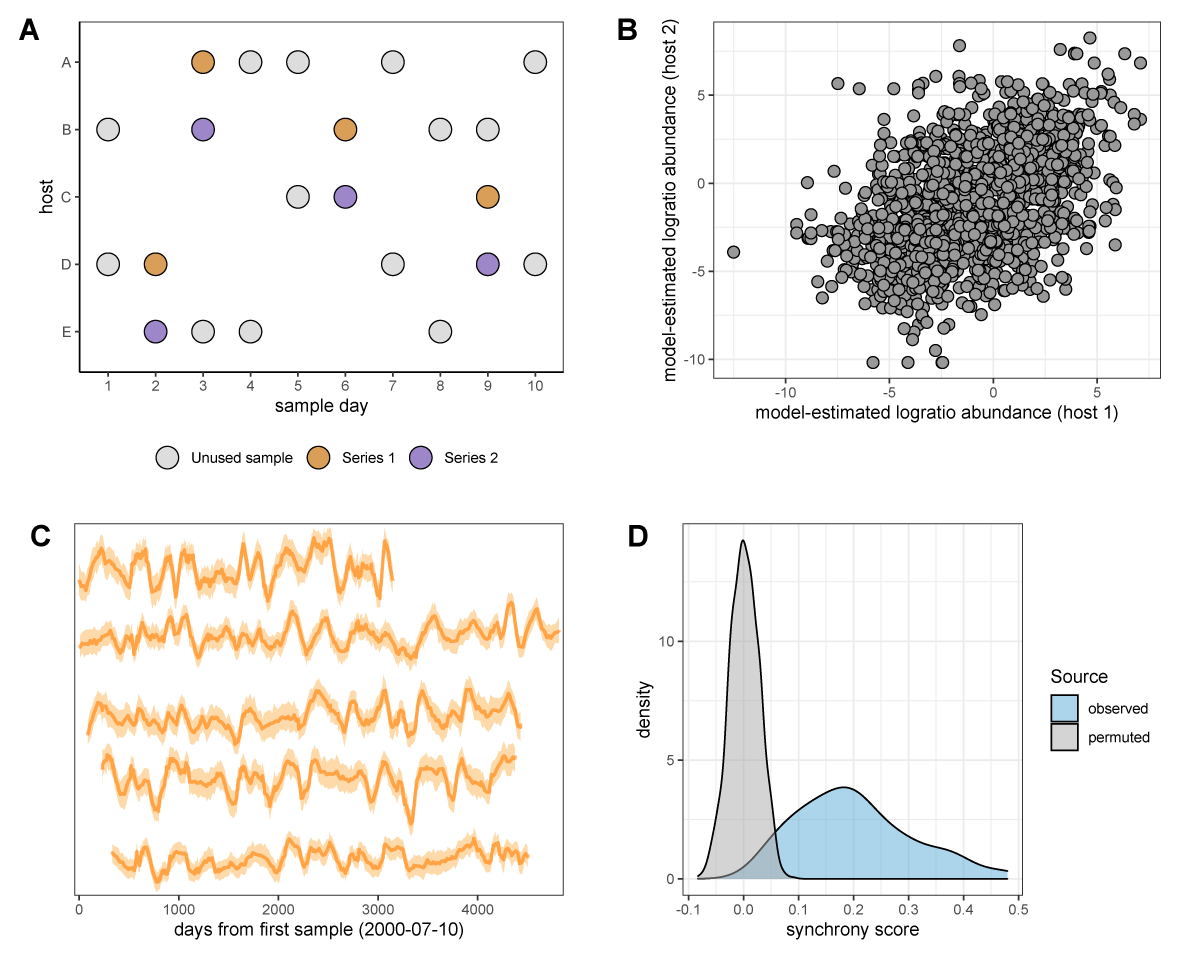

##### Figure S10. Measuring synchronized dynamics in the same ASV across many hosts

The cartoon in panel **(A)** shows the sample selection procedure. We selected pairs of samples from different hosts collected within 24 hours of each other, producing 20,000 “same-day” pairs (e.g., purple and mustard-colored pairs on days 2, 3, 6, and 9). We subset these pairs to one same-day pair per host pair, producing 1540 same-day pairs from thinned host series and extracted the correlated ASV abundance from these pairs. Synchrony for a given taxon is estimated as the correlation of the centered log-ratio abundance of that taxon across Series 1 and 2. Panel **(B)** shows the correlation between log-ratio abundance estimates in paired host samples for the most synchronous taxon, ASV #21 (Clostridiaceae 1; r = 0.480). Panel **(C)** shows the time-aligned model-estimated centered log-ratio abundances of ASV #23 in five well-sampled hosts that each lived in a different baboon social group (hosts from top to bottom are F09, F31, F35, F27, F20). Panel **(D)** shows the distribution of observed synchrony estimates for all 125 ASVs (light blue) compared to synchrony estimates from a permuted/null distribution (gray). 118 of 125 ASVs had significantly higher synchrony than expected (FDR ≤ 0.05; permutation scheme).

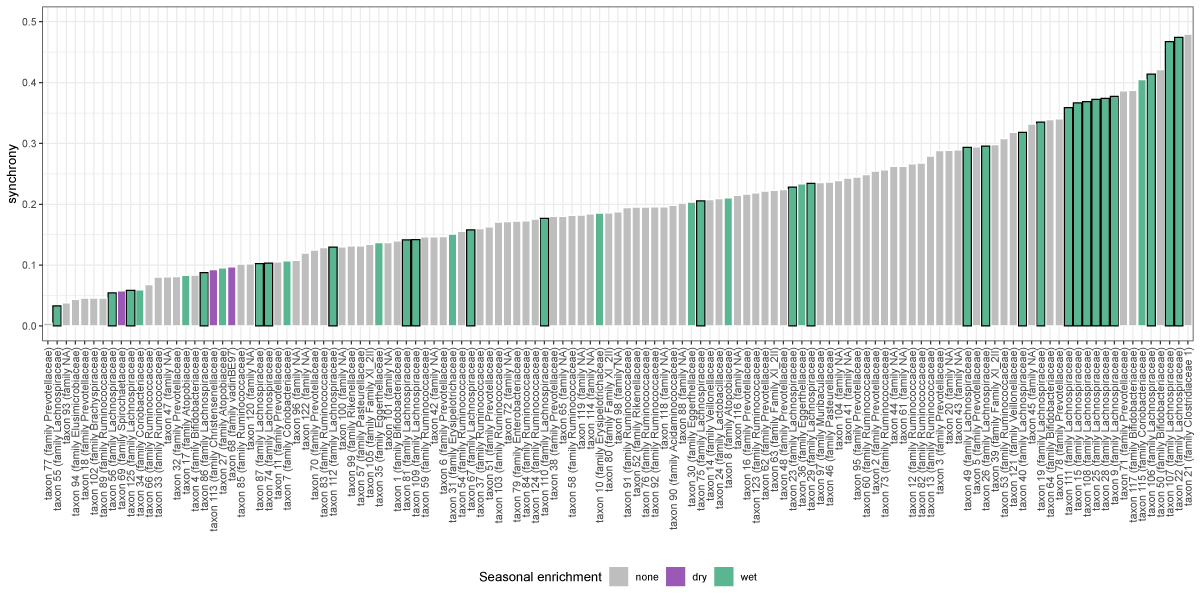

##### Figure S11. Seasonality is not associated with synchrony

Synchrony estimates for all ASVs are given as bar plots. ASVs that were identified as having seasonally differential CLR abundance in Björk et al. (2022) are labeled by the season in which they were more abundant: dry or wet. Seasonally variable labels do not predict synchrony (ANOVA, p=0.358).

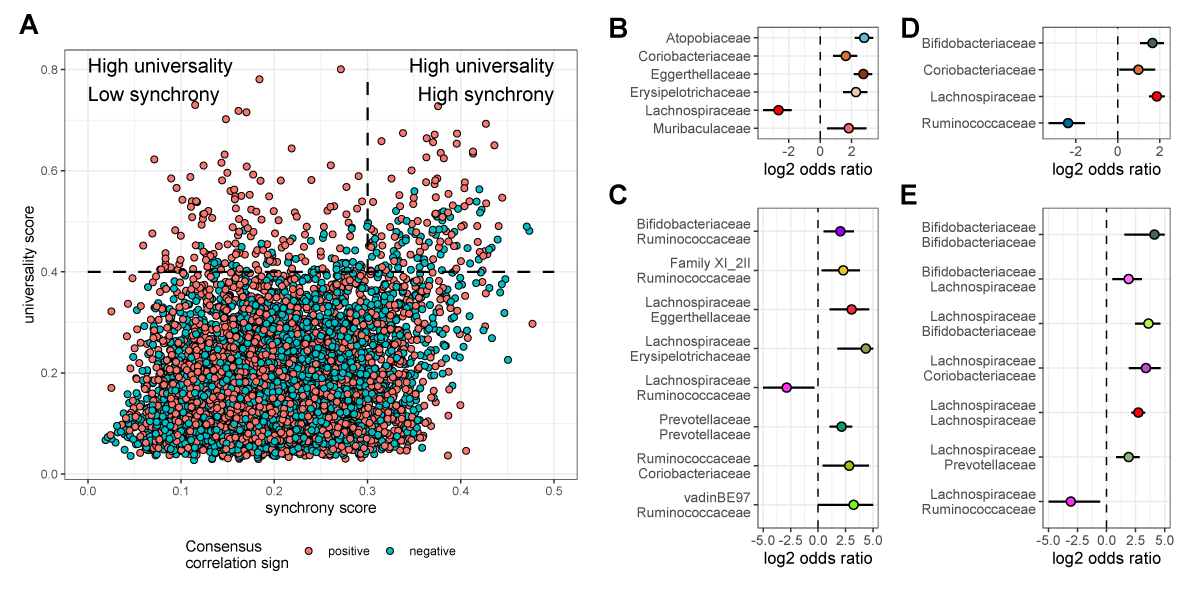

##### Figure S12. Synchrony weakly predicts universality

Panel **(A)**: mean ASV-ASV pair synchrony estimates are plotted against universality scores for all 7750 ASV pairs. Synchrony weakly predicts universality (r=0.264, p < 0.0001). Panel **(B)**: ASV-ASV pairs with low synchrony and high universality scores (i.e., top-left corner of scatter plot in panel **(A)**) are relative enriched for members of the family Atopobiaceae (p < 0.0001; **Table S9**) while pairs with high observed synchrony and high universality scores are enriched for members of the family Lachnospiraceae (p < 0.0001; **Table S9**). Panels (**C**) and (**D**) likewise show family-family pair enrichment in high universality, low synchrony or high universality, high synchrony pairs.

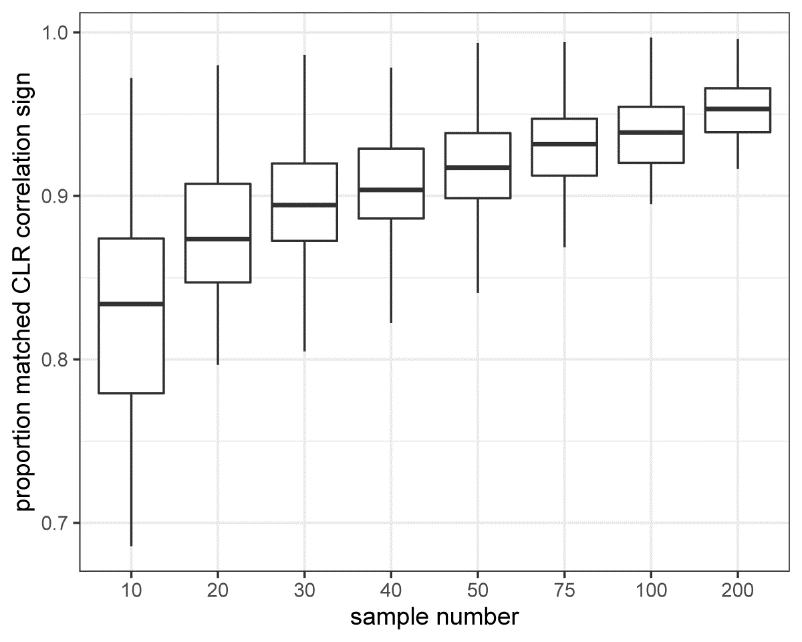

##### Figure S13. Informative sample number

Proportion agreement in sign between elements of simulated and estimated CLR taxon-taxon correlation matrices as a function of down-sampled sample number. Data were simulated from the *basset* model, randomly downsampled to between 10 and 200 samples, and correlation between log ratio taxa was estimated on this subset of samples by *basset*. A per-host minimum sample set size of at least 75 was selected on this basis: at an average sample size of 75, well over 90% of elements in the estimated covariance matrix were inferred with the correct sign on average.

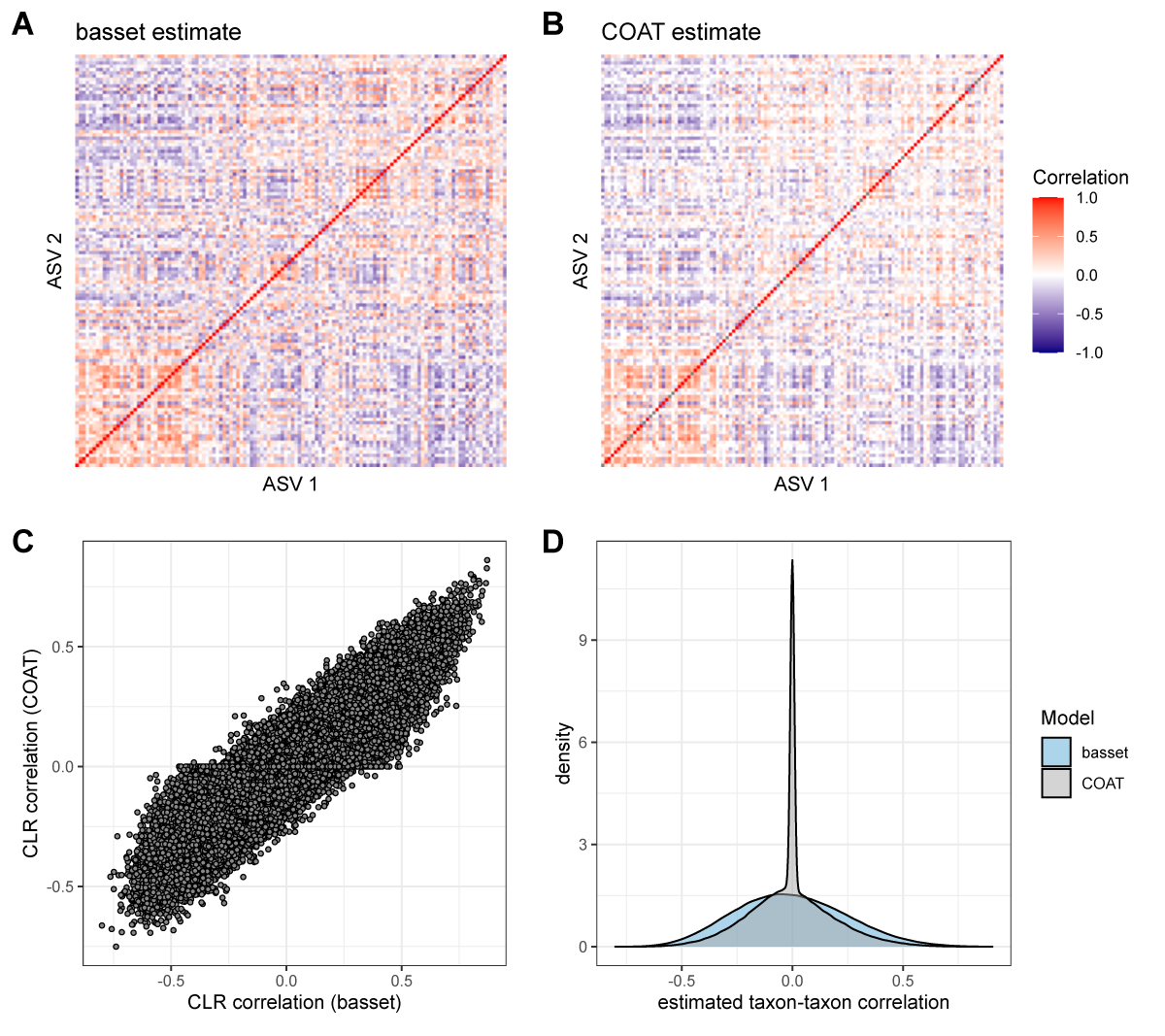

##### Figure S14. Concordance with COAT estimates

COAT (Cao et al., 2021) estimates regularized CLR correlation from independent samples. Panels **(A)** and **(B)** show estimated CLR ASV-ASV correlation matrices for samples from host F26 from *basset* and COAT, respectively. Panel **(C)** shows COAT vs. *basset* estimates of ASV-ASV correlation over all hosts and ASV-ASV pairs (r=0.884). COAT’s strong shrinkage of estimates is apparent in panel **(D)**.

### References

1.	Björk, J., et al., *Synchrony and idiosyncrasy in the gut microbiome of wild primates.* Nature Ecology & Evolution, 2022. **6**: p. 955-964.

2.	Grieneisen, L., et al., *Gut microbiome heritability is near-universal but environmentally contingent.* Science, 2021. **373**: p. 181-186.

3.	Cao, Y., W. Lin, and H. Li, *Large covariance estimation for compositional data via composition-adjusted thresholding. .* J Am Stat Assoc, 2019: p. 759-772.
